# Examining dynamics of three-dimensional genome organization with multi-task matrix factorization

**DOI:** 10.1101/2023.08.25.554883

**Authors:** Da-Inn Lee, Sushmita Roy

## Abstract

Three-dimensional (3D) genome organization, which determines how the DNA is packaged inside the nucleus, has emerged as a key component of the gene regulation machinery. High-throughput chromosome conformation datasets, such as Hi-C, have become available across multiple conditions and timepoints, offering a unique opportunity to examine changes in 3D genome organization and link them to phenotypic changes in normal and diseases processes. However, systematic detection of higher-order structural changes across multiple Hi-C datasets remains a major challenge. Existing computational methods either do not model higher-order structural units or cannot model dynamics across more than two conditions of interest. We address these limitations with Tree-Guided Integrated Factorization (TGIF), a generalizable multi-task Non-negative Matrix Factorization (NMF) approach that can be applied to time series or hierarchically related biological conditions. TGIF can identify large-scale changes at compartment or subcompartment levels, as well as local changes at boundaries of topologically associated domains (TADs). Compared to existing methods, TGIF boundaries are more enriched in CTCF and reproducible across biological replicates, normalization methods, depths, and resolutions. Application to three multi-sample mammalian datasets shows TGIF can detect differential regions at compartment, subcompartment, and boundary levels that are associated with significant changes in regulatory signals and gene expression enriched in tissue-specific processes. Finally, we leverage TGIF boundaries to prioritize sequence variants for multiple phenotypes from the NHGRI GWAS catalog. Taken together, TGIF is a flexible tool to examine 3D genome organization dynamics across disease and developmental processes.

## Introduction

The three-dimensional (3D) organization of the genome refers to the packaging of DNA inside the nucleus. It has emerged as a key regulatory mechanism of cellular function and dysfunction across diverse developmental (Zheng and Xie, 2019), disease (Lupiáñez et al., 2016), and evolutionary contexts (McCord, 2017; Eres et al., 2019). High-throughput chromosomal conformation capture (Hi-C) technologies enable the study of 3D genome organization by experimentally measuring the tendency of genomic regions to spatially interact with one another (Kempfer and Pombo, 2020; Mumbach et al., 2016; Kempfer and Pombo, 2019; Dekker et al., 2023). The 3D genome is organized into structural units at multiple scales: compartments spanning several megabases, Topologically Associated Domains (TADs) spanning hundreds of kilobases scale, and enhancer-promoter loops involving pairs of loci of a few thousand bases. (Bouwman and de Laat, 2015; Rowley and Corces, 2018; Kempfer and Pombo, 2020). Changes in 3D genome organization at different topological levels have been observed with transitions in both normal (Bonev et al., 2017; Stadhouders et al., 2018; Zheng and Xie, 2019) and disease processes (Lupiáñez et al., 2016; Norton and Phillips-Cremins, 2017; Wang et al., 2021). For example, during differentiation of mouse embryonic stem cells to a neuronal lineage, changes in topological structure are associated with cell fate specification and gene expression changes (Bonev et al., 2017). Changes in 3D genome organization have also been seen in immune response to viral infections (Wang et al., 2021) and diseases such as cancer (Hnisz et al., 2016; Akdemir et al., 2020; Dubois et al., 2022). Through efforts from large-scale consortia such as the 4D Nucleome project, Hi-C measurements are becoming increasingly common from multiple conditions corresponding to time points, cell types and species (Dekker et al., 2017; Reiff et al., 2022; Dekker et al., 2023; Roy et al., 2023). These datasets provide a unique opportunity to examine the dynamics of 3D genome organization across space and time and its impact on disease and normal processes.

Reliable detection of 3D genome dynamics at different units of organization is a significant computational challenge. Current computational approaches to examine dynamics in the 3D genome can be grouped into those that identify large-scale or compartmental-level changes (Fotuhi Siahpirani et al., 2016; Chakraborty et al., 2022), those that can identify TAD-scale changes or “differential TADs” (Wang et al., 2020; Cresswell and Dozmorov, 2020), and those that examine changes at the level of loops or interactions (Ardakany et al., 2019; Lun and Smyth, 2015; Djekidel et al., 2018; Galan et al., 2020; Stansfield et al., 2019). Compared to methods for detecting differences at the loop level, there are relatively few approaches to detect TAD or compartment changes. The most common approach to study TAD dynamics across multiple conditions is to first apply a TAD-calling method to data from each condition, followed by post-processing to identify TAD boundaries in one condition but not another (Zhang et al., 2019; Bonev et al., 2017; Stadhouders et al., 2018; Wang et al., 2022; Emerson et al., 2022). While such a two-step approach can identify some meaningful differences, the unsupervised nature of TAD finding could make these approaches more susceptible to finding non-biological differences. Numerous studies have shown that Hi-C count profiles obey cell type, timepoint and species relationships, where datasets from nearby contexts are more similar than those that are far away (Bonev et al., 2017; Zhang et al., 2019; Yang et al., 2017; Vietri Rudan et al., 2015). An approach that constrains the TAD and compartment finding based on such prior information about the relationships between the input datasets could be less prone to spurious differences. A few methods have been developed to directly identify TAD boundary differences, but they are focused on pairs of conditions (Wang et al., 2020) or limited in their ability to compare more than two conditions (Cresswell and Dozmorov, 2020).

To address the dearth of methods for identifying large-scale organizational changes, especially when considering more than two datasets, we developed Tree-Guided Integrated Factorization (TGIF), a multitask learning framework using Non-negative Matrix Factorization (NMF) to enable joint identification of organizational units such as compartments and TADs across multiple conditions. NMF is a popular dimensionality reduction approach that has been used for analyzing genomic data (Stein-O’Brien et al., 2018; Kotliar et al., 2019; Lee and Roy, 2021) as well as images (Lee and Seung, 1999; Kalayeh et al., 2014), where the low-dimensional factors can recover the major patterns in the data. In the case of Hi-C data, the output factors represent the lower-dimensional view of the chromosomal architecture. TGIF can take as input multiple Hi-C matrices from related biological conditions, for example, different time points, treatments, diseases or cell types. TGIF uses hierarchical multi-task learning to constrain the lower-dimensional factors from closely related tasks (e.g. consecutive time points) to be similar. We use the low dimensional factors for each of the conditions to identify changes at both the compartment and TAD levels.

We applied both TGIF-DB and TGIF-DC to three different mammalian differentiation timecourse Hi-C datasets: a 2 timepoint dataset comprising human pluripotent cell line H1 and differentiated endoderm (Reiff et al., 2022; Dekker et al., 2023), a 3 timepoint dataset of mouse neural differentiation (Bonev et al., 2017), and a 6 timepoint dataset collected during human cardiomyocyte differentiation (Zhang et al., 2019). Compared to existing approaches, TGIF-DB TAD boundaries are more enriched in CTCF binding and is less susceptible to false differential boundaries that could arise due to non-biological factors such as different replicates, different downsampling depths, normalization methods, and binning resolutions.

At the compartment level, TGIF-DC differential compartmental regions displays significant change in accessibility and gene expression, while recovering compartments of similar quality compared to competing methods, measured by chromatin accessibility and observed-over-expected interaction counts. TGIF-DC can additionally cluster genomic regions into the more granular subcompartments with distinct histone modification patterns. We used TGIF-DB boundaries to assess the extent to which differentially expressed genes localize near significantly differential boundaries. Finally, we use TGIF boundaries identified in cardiomyocyte differentiation data to interpret sequence variants identified from the NHGRI Genome Wide Association Studies (GWAS) catalog and identify cardiovascular disease associated SNPs to be enriched in TGIF boundaries. Together, these results demonstrate the versatility and utility of TGIF to examine changes in higher-order 3D genome organization across diverse types of dynamic processes.

## Results

### Tree-guided Integrated Factorization (TGIF) for examining dynamics in 3D genome organization

Tree-guided Integrated Factorization (TGIF) is a general-purpose framework to study 3D genome organization dynamics both at the TAD and compartment levels (**Figure 1**). TGIF is based on multi-task non-negative matrix factorization (NMF). It takes as input a set of Hi-C matrices, each representing a biological condition, and a user-specified tree structure that can encode an arbitrary relationship among the conditions, such as time or cell type lineage (**Figure 1**, **Supp Figure 1**). TGIF uses a novel regularization term in its objective to jointly factorize the matrices such that input matrices from more closely related conditions result in more similar lower-dimensional representations, i.e., factors.

**Figure 1.**
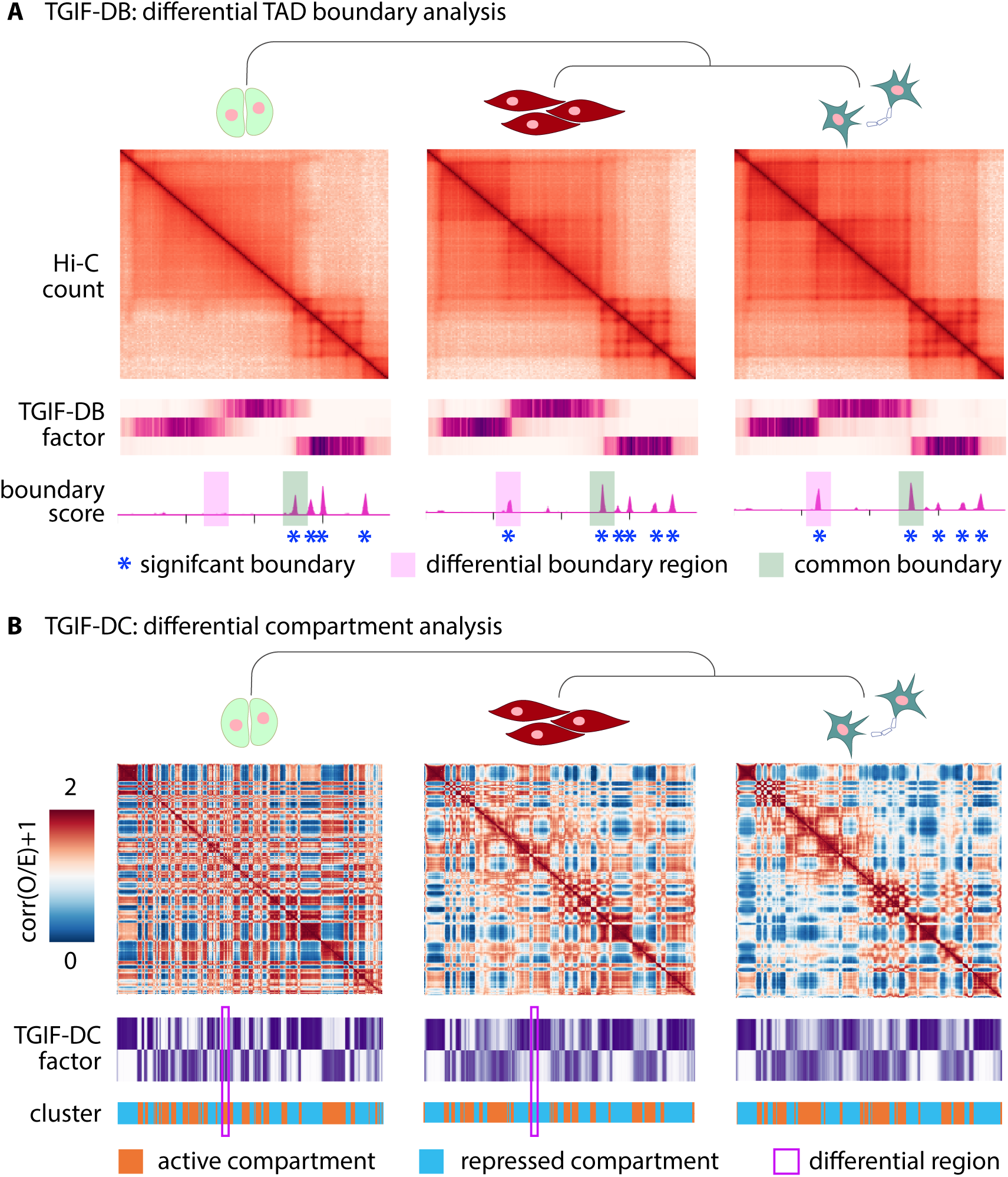
Overview of TGIF. **(A)** TGIF for differential boundary analysis (TGIF-DB). TGIF-DB takes multiple Hi-C count matrices as input and simultaneously learns a lower dimensional representation of genomic regions based on their interaction patterns. The input matrices are from related biological conditions with their relationship encoded as a tree. From the lower-dimensional factors, we measure the boundary score of each region, identify boundaries for each input condition and significantly differential boundaries for every pair of conditions. **(B)** TGIF for differential compartment analysis (TGIF-DC). TGIF-DC converts input matrices into correlation matrices of observed-over-expected (O/E) counts and factorizes them to yield latent features, which are used to cluster the regions. Each cluster correspond to a compartment or a subcompartment. TGIF-DC also identifies significantly differential compartmental regions for every pairs of input conditions.

To handle both compartment and TAD identification, we implemented two versions of TGIF: TGIF-DB and TGIF-DC. TGIF-DB identifies conserved and differential boundaries demarcating TADs under different conditions (**Figure 1A**, **Supp Figure 1A**, **Methods**), while TGIF-DC identifies compartment-level changes in 3D genome organization (**Figure 1B**). In TGIF-DB, the factorization is performed on sub-matrices along the diagonal of the intrachromosomal Hi-C matrices, as these diagonal sub-matrices capture the TAD-scale, local topology of chromosomes. Each sub-matrix is factorized over a range of *k*, the hyper-parameter specifying the rank of the lower-dimensional space (**Supp Figure 1B**). TGIF-DB calculates a boundary score from the factors at each *k*, which are averaged to provide an overall boundary score (**Supp Figure 1C**). Considering multiple *k* allows us to capture structural units or domains of different sizes in the lower dimensional space and removes the need to specify the number of factors (**Methods**). TGIF-DB identifies regions with significant boundary scores by comparing the average boundary scores against a “null distribution” of boundary scores to calculate an empirical p-value (**Supp Figure 1D**). TGIF-DB outputs the list of significant boundaries corresponding to each input dataset and a list of significantly differential boundary regions for every pair of input count matrices (**Supp Figure 1E**, **Methods**). A number of optional outputs that can be used for additional downstream analysis (See **Availability** section for more details).

TGIF-DC operates at the entire chromosome level and applies its multi-task factorization on the observed-over-expected (O/E) counts matrix as described previously (Lieberman-Aiden et al., 2009; Rao et al., 2014, **Methods**). To identify the two major compartments of active and repressive genomic regions, TGIF-DC factorizes the O/E matrices with parameter *k* = 2. The resulting factors are used to group the genomic regions into 2 different clusters. By specifying a higher parameter value, e.g. *k* = 5, TGIF-DC can also identify more granular subcompartment structures, which can be interpreted using one-dimensional chromatin signals. Similar to TGIF-DB, TGIF-DC identifies significantly differential compartment and subcompartment regions for every pair of input conditions (**Methods**).

In cases where the relationship between the input Hi-C data is not available (e.g. integrating Hi-C datasets from multiple studies or pseudo-bulk single-cell Hi-C data from cell clusters; Zhou et al., 2019; Zhang et al., 2022) TGIF can infer a tree structure based on the pairwise similarity of the input Hi-C matrices measured by stratum-adjusted correlation coefficient (SCC; Yang et al., 2017, **Methods**, **Supp Figure 2**) or a similar distance-stratified metric.

### TGIF-DB identifies CTCF-enriched boundaries and fewer false-positive differential boundaries

TGIF-DB was benchmarked against four other TAD calling methods: three methods designed for calling TADs and boundaries from a single Hi-C matrix (which we refer to as single-task methods), and one designed specifically for differential boundary identification (**Methods**). The three single-task methods were: (1) GRINCH (Lee and Roy, 2021), which uses NMF for TAD identification; (2) SpectralTAD (Cresswell et al., 2020), which also uses dimension reduction; and (3) TopDom(Shin et al., 2016), which uses changes in average contact frequencies upstream and downstream of a given genomic region to determine significant boundaries. TADCompare (Cresswell and Dozmorov, 2020) is a method designed for differential boundary detection and takes as input pairs of input Hi-C matrices and identifies non-differential and differential boundaries between them. Default or recommended parameters were used for all benchmarking experiments (**Methods**).

Since real Hi-C datasets do not have ground-truth set of TAD boundaries, we first evaluated the quality of TAD boundaries identified by each method based on the enrichment of CTCF binding. CTCF is an architectural protein associated with establishing boundaries (Merkenschlager and Nora, 2016; Gómez-Díaz and Corces, 2014; Cubeñas-Potts and Corces, 2015). We used the time-series dataset of cardiomyocyte differentiation (Zhang et al., 2019) which profiled both genome-wide chromosome conformation with Hi-C and CTCF binding with ChIPseq. The single-task methods, GRiNCH, SpectralTAD, and TopDom, were applied to each of the six timepoints (day 0, 2, 5, 7, 15, 80) from the cardiomyocyte dataset independently. For TADCompare, we gave the algorithm pairs of consecutive timepoints (day 0 vs 2, 2 vs 5, 5 vs 7, 7 vs 15, 15 vs 80) to find non-differential and differential boundaries between each pair of timepoints (**Methods**). Finally, TGIF-DB was applied to all available timepoints together with an input tree structure capturing the temporal dependency (see **Supp Figure 3** for input tree). The resulting boundaries for each timepoint was used for CTCF enrichment analysis. Fold enrichment of CTCF peaks in the boundary regions was calculated against the genomic background (**Methods**). We find that significant boundaries identified by TGIF-DB has the highest fold enrichment, followed by single-task methods, TopDom and GRiNCH (**Figure 2A**).

**Figure 2.**
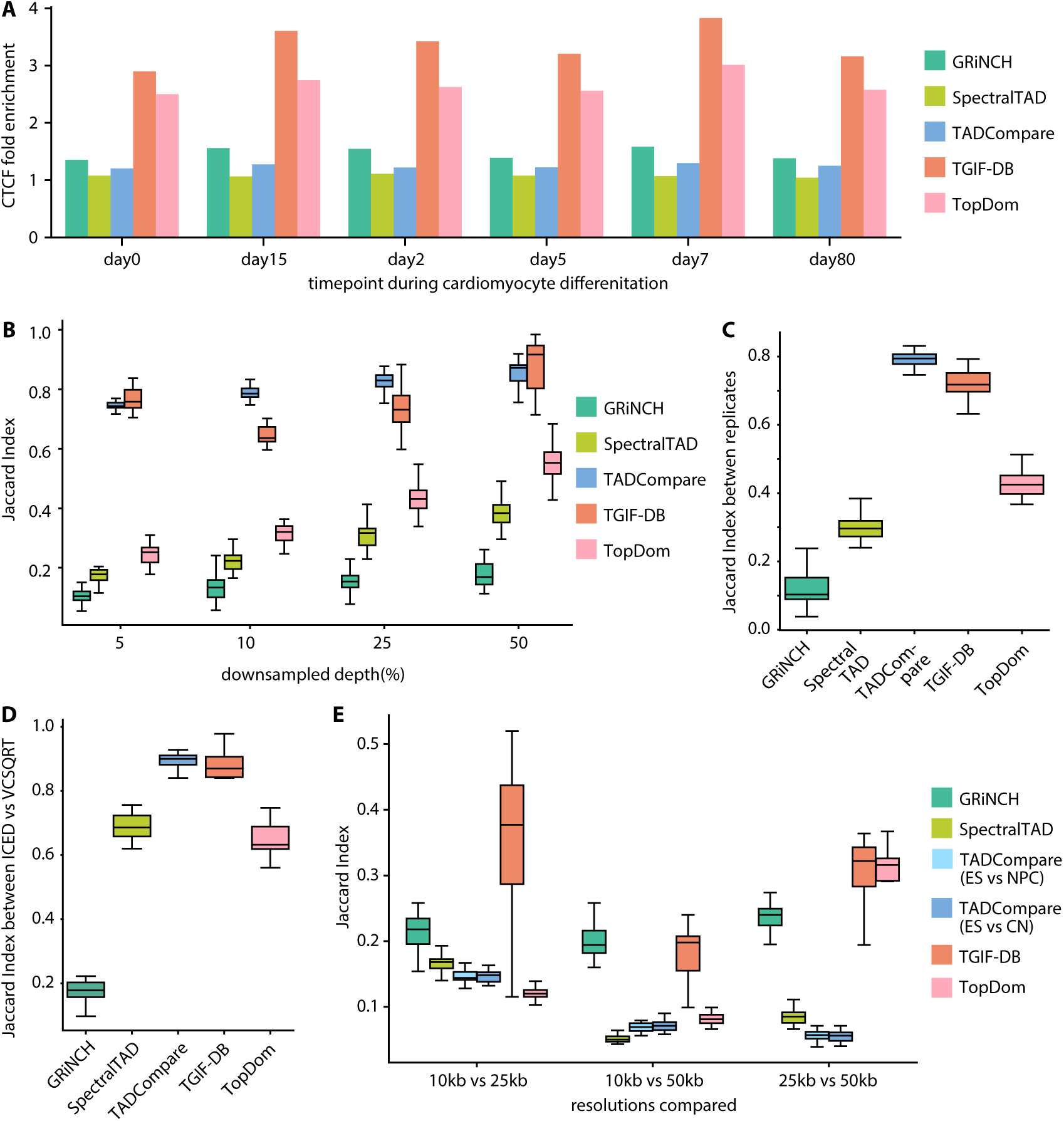
Benchmarking TGIF-DB. **(A)** CTCF peak enrichment in boundaries from different TAD- calling and differential-boundary-calling methods. **(B)** Boundary set similarity measured by Jaccard index between GM12878 data and downsampled data, across different downsampling depths. **(C)** Boundary set similarity between biological replicates of hESC (from day 0 of cardiomyocyte differentiation data). **(D)** Boundary set similarity between ICE-normalized and VCSQRT-normalized input matrices of mESC (from mouse neural differentiation data). **(E)** Similarity of boundary sets from input data of different resolutions (10kb, 25kb, and 50kb).

We next benchmarked the ability to detect true versus false boundary differences between Hi-C datasets, which could arise due to various non-biological reasons such as: (1) datasets with different depths, (2) datasets from biological replicates, (3) different normalization process, (4) different bin resolution. Since the underlying biological process is the same, any differences in boundaries between such datasets are considered as false positives. We took a high-depth Hi-C dataset from the GM12878 cell line with 4.01 billion reads in total (Rao et al., 2014; Reiff et al., 2022) and subsampled 5, 10, 25, 50% of the reads to create downsampled versions of the data (**Methods**). As these datasets represent the same cell line, any differential boundaries can be considered as false positives resulting from the depth difference. We again applied the single-task methods (GRiNCH, SpectralTAD, TopDom) to each of the original and downsampled datasets independently. Differential-TAD methods (TADCompare and TGIF-DB) were applied to a pair of datasets, i.e. the original high-depth and a downsampled. We measured the Jaccard index between the boundaries identified in a pair; the higher the Jaccard index, the fewer the false-positive differences identified by a method. Across all downsampled depths, TADCompare and TGIF-DB were the top performing methods with consistently high Jaccard scores; TADCompare outperformed TGIF-DB at 10 and 25% depth, and TGIF-DB outperformed TADCompare at 50% (**Figure 2B**). Single-task methods (GRiNCH, SpectralTAD, and TopDom) had much lower Jaccard score with discrepancy increasing with depth differences.

We next compared methods for their ability to recapitulate TAD structure across biological replicates (**Methods**) by applying them to the day 0 biological replicates of the cardiomyocyte differentiation dataset (Zhang et al., 2019), which represents the H1 ESC state. This time point is expected to have the lowest artificial differences as it is from a cell line and the process of differentiation can introduce additional sources of heterogeneity. We measured the Jaccard coefficient between the boundaries identified in the pair of replicates. Here again TADCompare and TGIF-DB had the highest Jaccard coefficients recovering most similar set of boundaries across the biological replicates (**Figure 2C**).

We also compared the methods for their ability to recover reproducible TADs across different normalization methods: Iterative Correction and Eigenvector decomposition (ICED, Imakaev et al., 2012) and square root vanilla coverage (VCSQRT, Rao et al., 2014) on a mouse embryonic stem cell (mESC) Hi-C dataset (Bonev et al., 2017, **Methods**). We observed similar results with TADCompare and TGIF-DB obtaining the highest Jaccard coefficients between TAD boundaries across different normalizations (**Figure 2D**).

Finally, we measured the stability of TAD boundaries to the changing resolution (10kb, 25kb, 50kb) of input Hi-C matrices using Jaccard index (**Methods**). TGIF-DB and GRiNCH yield the most stable or similar boundaries to changing resolution (**Figure 2E**), with the exception of 25kb-50kb where TopDom also performed well. Since both TGIF-DB and GRiNCH are based on non-negative matrix factorization, it is possible that NMF-based methods are less susceptible to changes in resolution of the dataset.

Taken together, these results demonstrate the advantages of using a multi-task matrix factorization method such as TGIF-DB to identify biologically relevant boundaries enriched in known boundary elements while minimizing false positive differences.

### TGIF-DC identifies compartment dynamics that are significantly enriched for differential regulatory signals

We compared the TGIF-DC compartments and differential compartments to three existing methods on intra-chromosomal count matrices at 100kb resolution from the H1 hESC and endoderm differentiation dataset. Two were single-task compartment calling methods: PCA-based (Lieberman-Aiden et al., 2009) and Cscore (Zheng and Zheng, 2018). The third method, dcHiC (Chakraborty et al., 2022), is a differential compartment identification method, and was applied in a pairwise comparison mode between H1 hESC and endoderm differentiated from H1 (**Methods**). TGIF-DC was applied to a simple tree with two leaf nodes as H1 and endoderm (**Methods**, **Supp Figure 3**).

We first compared the similarity of compartment assignments in H1 hESC from different methods by treating the assignment as clusters and using Rand Index (**Figure 3A**). The PCA-based method and dcHiC, which also utilizes PCA, produced the most similar compartments as expected (Rand Index: 0.91), followed by TGIF-DC (Rand Index: 0.79-0.8). Cscore found a substantially different set of compartments (Rand Index: 0.52). We next assessed the quality of compartments by measuring three cluster quality metrics, Silhouette Index (SI, **Figure 3B**), Calinski-Harabasz score (CH, **Figure 3C**), and Davies-Bouldin Index (DBI, **Figure 3D**), using observed-over-expected (O/E) counts as features of each genomic loci (**Methods**). In all three metrics, TGIF-DC, dcHiC and PCA-based compartments are comparable in their quality and outperformed Cscore.

**Figure 3.**
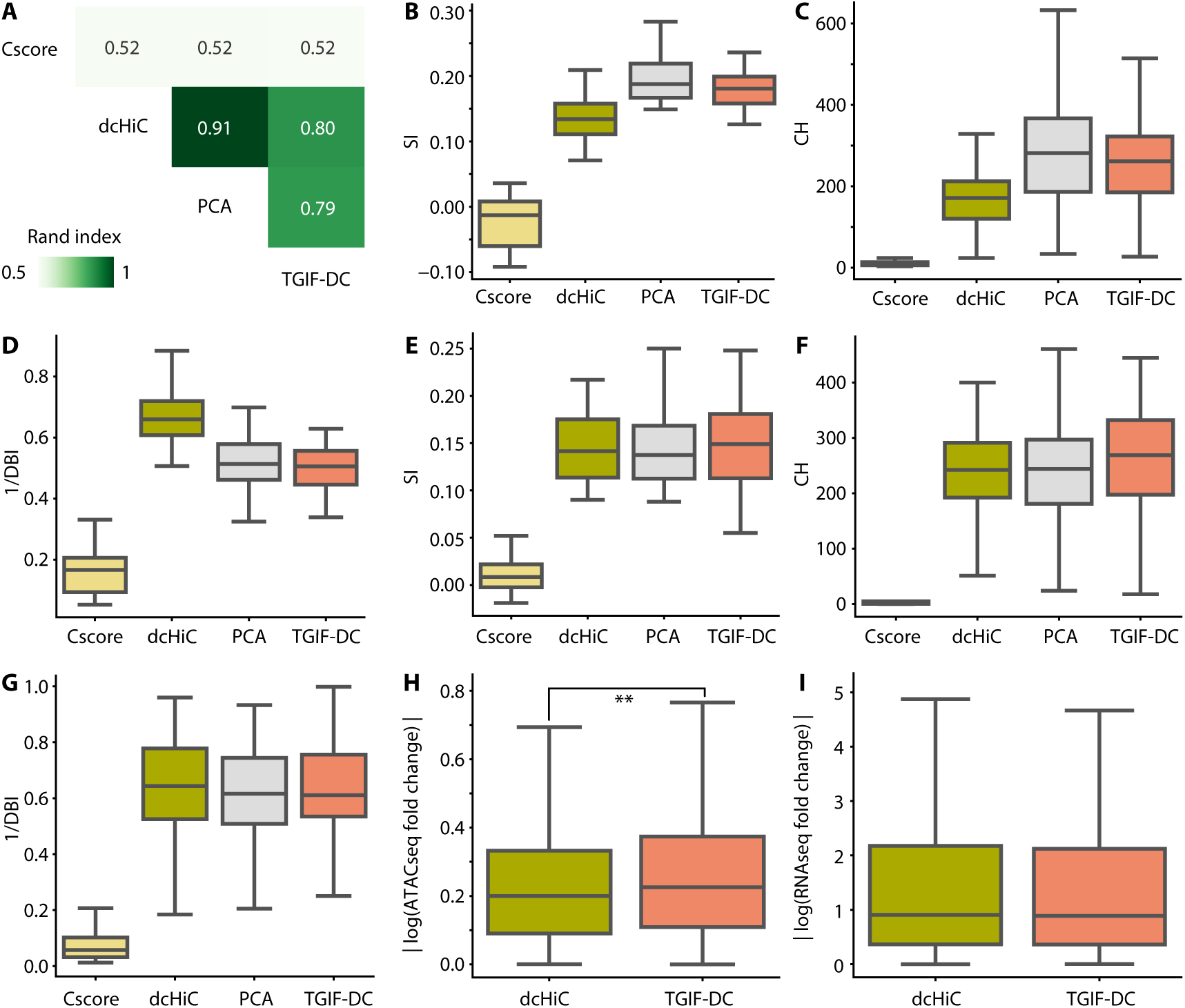
Benchmarking TGIF-DC on data from H1 and H1 differentiated to definitive endoderm. **(A)** Similarity of compartment assignments from different methods measured by Rand Index. **(B)** Quality of compartments based on O/E counts measured by Silhouette Index (SI), **(C)** Calinski-Harabasz score (CH), **(D)** Davies-Bouldin index (DBI). **(E)** SI, **(F)** CH, **(G)** DBI on accessibility (ATACseq) signal. **(H)** Magnitude of log fold change in accessibility between H1 and endoderm within significantly differential compartmental regions (sigDC) identified by dcHiC and TGIF-DC. **(I)** Magnitude of log fold change in gene expression between H1 and endoderm within sigDC identified by dcHiC and TGIF-DC.

We also assessed the compartment quality using chromatin accessibility, a key regulatory measurement that characterizes different compartment types (e.g., the active A and repressive B; Lieberman-Aiden et al., 2009; Fortin and Hansen, 2015), as the feature. Briefly, we used SI (**Figure 3E**), CH (**Figure 3F**) and DBI (**Figure 3G**) using the mean basepair ATACseq signal for each 100kb region (**Methods**). For all three metrics, the compartments from TGIF-DC, PCA-based method, and dcHiC are of similar quality and higher than Cscore.

Finally, we compared TGIF-DC exclusively with dcHiC, the only method among the three compared, that identifies *differential* compartment regions. Significantly differential compartmental regions (sigDC) identified by TGIF-DC have significantly higher change in accessibility signal and gene expression compared to regions not part of sigDC (**Supp Figure 4A,B**). Compared to significantly differential regions identified by dcHiC, sigDC regions from TGIF-DC also have significantly higher change in accessibility signal (*t*-test p-value *<*1e-2, **Figure 3H**) and are comparable in terms of the change in gene expression levels (**Figure 3I**).

### TGIF-DC offers a unified framework to identify both compartment and subcompartment dynamics

While compartments provide a global partitioning of each chromosome, the genome is hierarchically organized with compartments further partitioned into smaller subcompartments that could represent functionally distinct set of regions (Rao et al., 2014; Xiong and Ma, 2019). Unlike existing compartment finding methods that need additional clustering steps to define subcompartment structure, TGIF-DC has a tunable parameter (*k*, the rank of NMF factors) that can be used to identify subcompartments within the same framework. TGIF-DC’s low-rank dimensionality reduction framework lends itself naturally to identify this subcompartment structure. To demonstrate TGIF-DC’s ability to identify both compartments and subcompartments, we applied it to the mouse neural differentiation dataset with 3 timepoints: embryonic stem cell or ES, neural progenitors or NPC, and cortical neurons, CN (**Supp Figure 3,5,6**). This dataset additionally measured six different histone modification signals for NPC and CN that were beneficial for additional biological interpretation of TGIF-DC results (**Figure 4A**, **Methods**). We first analyzed the compartment structure from TGIF-DC (*k* = 2) for each chromosome, based on GC content (mean GC percentage for each 100kb bin, **Methods**), annotating the compartment with higher GC content as compartment A and the one with lower GC content as compartment B (**Methods**, **Supp Figure 7**). Regions annotated as A compartment by TGIF-DC have significantly higher signal for marks associated with active enhancer (H3K27ac, H3K4me1) or elongation (H3K36me3) than those in B compartment (**Figure 4B**).

**Figure 4.**
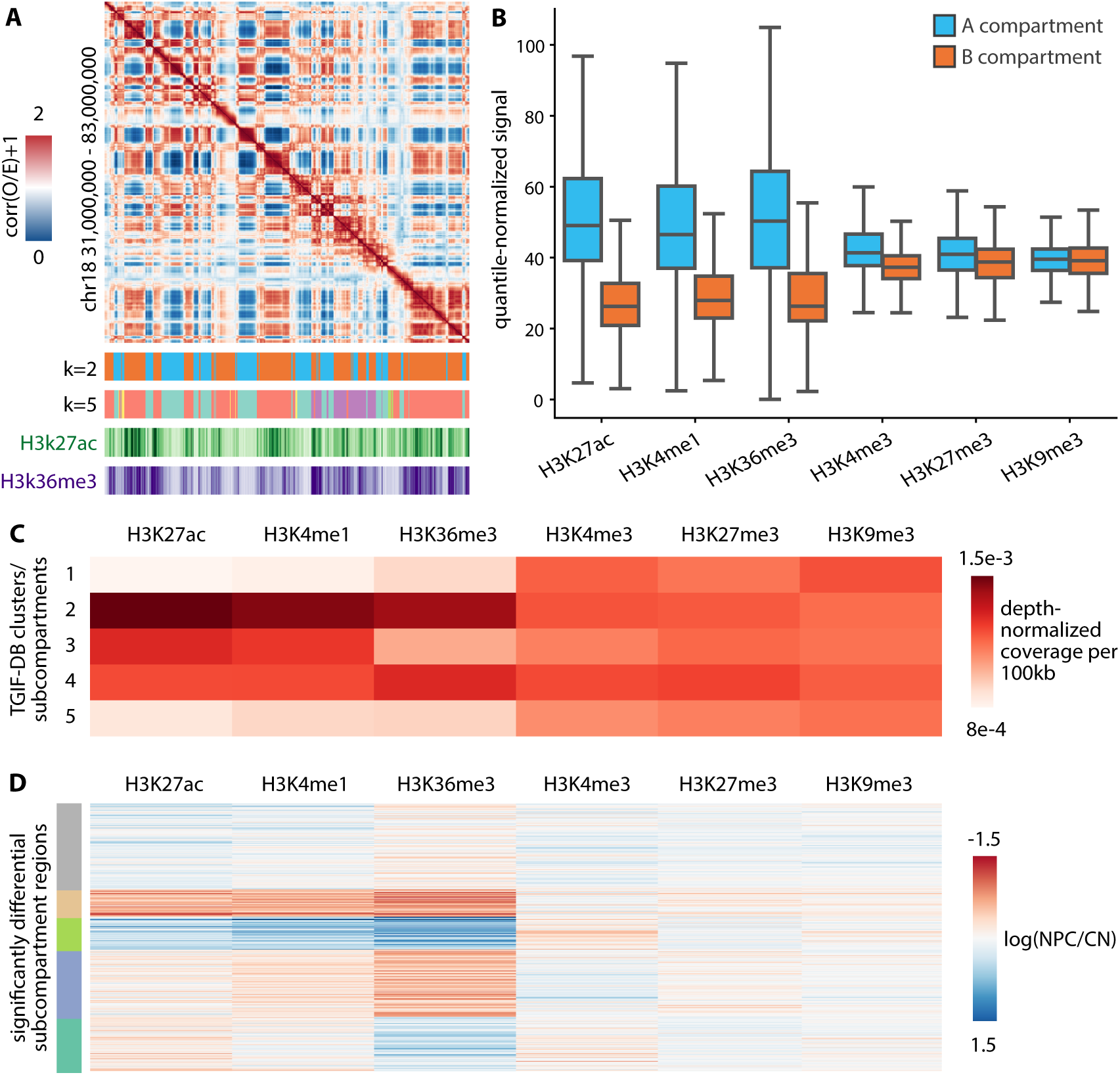
Characterizing compartments and subcompartments identified by TGIF-DC in mouse neural differentiation data. **(A)** A heatmap visualization of correlation matrix of O/E counts from cortical neuron (CN) chr18 regions at 100kb resolution, followed by TGIF compartment assignments (i.e. clusters from *k* = 2) and subcompartments (e.g. clusters from *k* = 5), and H3K27ac/H3K36me3 ChIPseq signal heatmaps. **(B)** Distribution of histone modification signal in A and B compartments in neural progenitors (NPC) and CN. **(C)** Mean histone modification signals across different subcompartments in NPC and CN. **(D)** Log fold change of histone modification signals between NPC and CN within significantly differential subcompartment regions identified by TGIF-DC. These regions were grouped based on their histone modification signal fold change patterns using k-means clustering and visualized here.

We next applied TGIF-DC with *k* = 5 to identify subcompartment structure per chromosome, each *k* corresponding to a different subcompartments (**Methods**). We interpreted these subcompartments based on the mean histone modification signal of the genomic loci assigned to each subcompartment. The subcompartments exhibited distinct histone modification patterns (**Figure 4C**, chr18), with subcompartments 1 and 5 associated with repressive marks (H3K9me3, H3K27me3), while the other three (2, 3 and 4) associated with active marks. Within these two groups, each subcompartment had a different signature of marks. For example, subcompartment 3 exhibits relatively lower signal of H3K36me3 compared to 2 and 4, while subcompartment 2 had a higher signal of all three activating marks (H3K27ac, H3K36me3, H3K4me1) compared to 3 and 4. Between the two subcompartments, 1 and 5, with repressive mark association, one (1) exhibited higher H3K4me3 and H3K9me3 levels compared to the other one (5).

Finally, we assessed TGIF-DC’s differential subcompartments by measuring the log fold change in histone modification signals between two timepoints, NPC and CN, and k-means clustering the regions based on this signal difference. We find distinct subgroups of regions with different fold change of the three activating marks, H3K27ac, H3K36me3, and H3K4me1 (**Figure 4D**). Interestingly, the repressive marks or the promoter specific mark, H3K4me3, did not vary substantially for these regions.

Taken together, these results demonstrate TGIF-DC’s flexible framework to identify both compartment and subcompartment level dynamics that are are associated with significant changes in regulatory activity between the timepoints or cell stages compared.

### Changes in gene expression are associated with changes in boundaries during differentiation

Untangling the relationship between 3D genome organization and gene expression remains a key question in regulatory genomics. While a direct mechanistic link between transcription and 3D genome organization has been observed (van Steensel and Furlong, 2019; Heinz et al., 2018) during cell state transitions (Pollex et al., 2024; Chen et al., 2024), other studies did not reveal changes in 3D genome organization to be a strong determinant of gene expression changes (Ing-Simmons et al., 2021; Espinola et al., 2021). To assess the extent to which changes in 3D genome structure are associated with changes in expression, we analyzed differential structural regions identified by TGIF-DC and TGIF-DB with differential gene expression in multiple mammalian differentiation datasets.

We applied TGIF-DC and TGIF-DB to the three different timecourse datasets with both Hi-C and RNAseq measurements (**Supp Figure 3**, **Supp Table 1-3**): (1) the two-timepoint dataset of H1 hESCs differentiated to endoderm state (Reiff et al., 2022; Dekker et al., 2023, **Supp Figure 4C-F**, **Supp Figure 8**), (2) the three-timepoint of mouse neural differentiation time course from mESC to cortical neurons (CN, Bonev et al., 2017, **Supp Figure 5,6,9**), and (3) the six-timepoint human cardiomyocyte differentiation timecourse from hESC to ventricular cardiomyocytes (Zhang et al., 2019, **Supp Figure 10,11**). For each time course, we performed pairwise comparison of differential boundary, compartment, and expression between all pairs of time points, e.g. H1 vs endoderm, mESC vs NPC, day 0 vs day 2 of cardiomyocyte differentiation. For each pair of timepoints, we performed a region-centric and gene-centric analysis to ask whether differentially expressed (DE) genes are enriched in three different sets of dynamic regions (**Methods**, **Figure 5A**): (A) regions near (i.e., within 100kb) of significantly differential boundaries (sigDB), (B) regions within a TAD with at least one sigDB, and (C) regions within significantly differential compartmental regions (sigDC). For the region-centric analysis, we measured the fold enrichment of regions overlapping a DE gene among dynamic regions compared to all genomic regions. For the gene-centric analysis, we measured the fold enrichment of DE genes among genes overlapping a dynamic region compared to the proportion of DE genes among all genes. Dynamic regions in set A (within 100kb of sigDB) are consistently enriched for DE genes across all datasets and timepoints compared (**Figure 5B** top, **Supp Table 5-7**). Furthermore, genes in set A are also enriched for DE genes compared to all genes (**Figure 5B** bottom). We did not see as much significant enrichment for regions in set B (within a TAD with at least one sigDB), likely because of the permissive inclusion criteria for set B. However, genes in set B were enriched for DE genes as well, although to a lower extent. When examining compartments, we found that regions in set C (within sigDC) are also significantly enriched in DE genes for the H1-endoderm differentiation and majority of the comparisons of the cardiomyocyte differentiation. The enrichment for genes was lower again due to the large number of genes within compartments.

**Figure 5.**
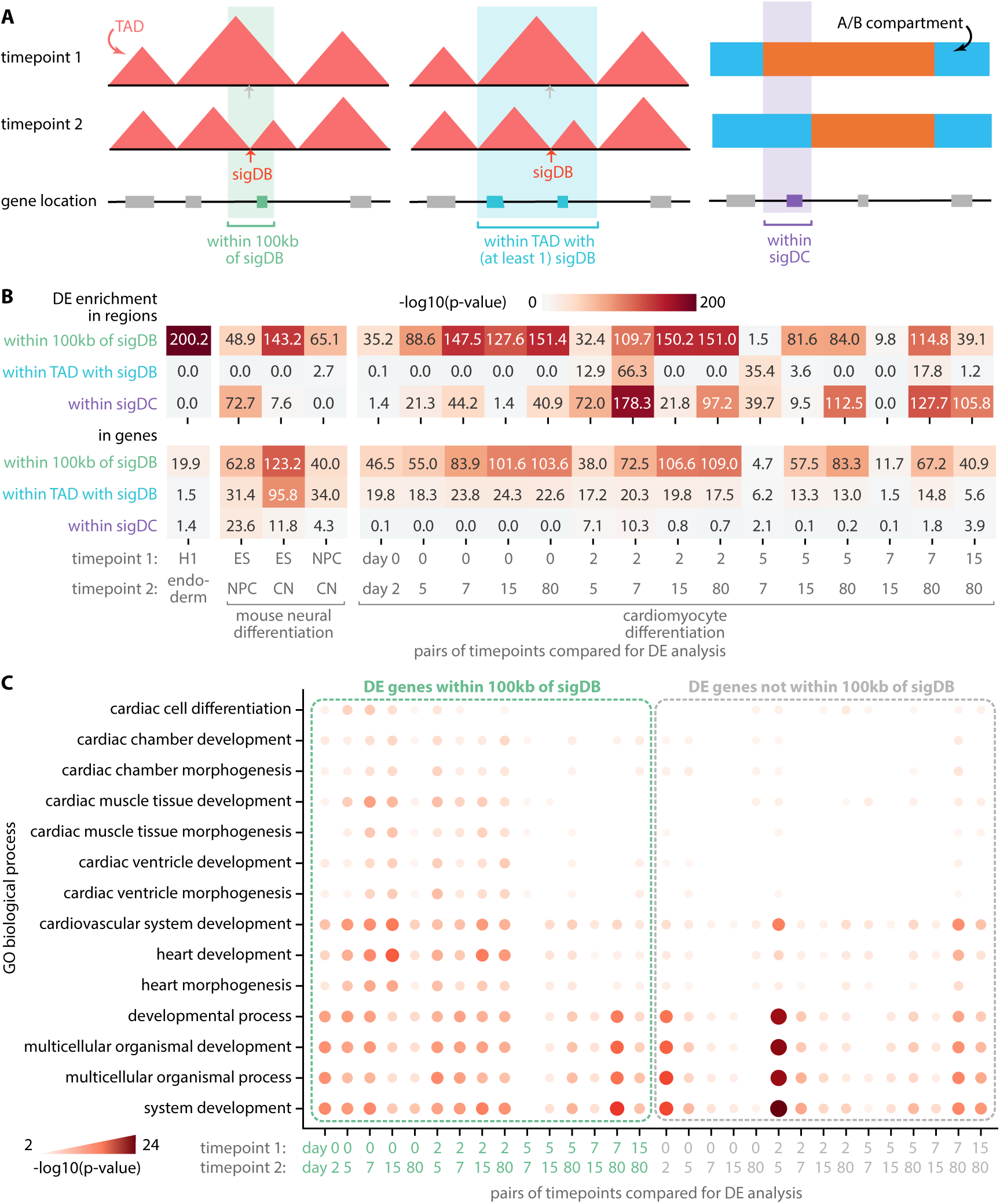
Differential gene expression near or within differential structural features. **(A)** Differential gene expression (DE) enrichment was measured in regions and genes near or within dynamic regions, i.e. regions within 100kb of significantly differential boundary (sigDB), regions within TAD with at least one sigDB, and regions within significantly differential compartmental regions (sigDC). **(B)** DE, sigDB, and sigDC were measured and identified in pairwise comparisons of timepoints across 3 mammalian differentiation datasets: H1 differentiated to endoderm, mouse neural differentiation (ES, NPC, CN), and cardiomyocyte differentiation (day 0, 2, 5, 7, 15, 80). Negative log p-value of the enrichment hypergeometric test is visualized here. **(C)** GO biological process enrichment of genes within 100kb of sigDB from cardiomyocyte differentiation data.

To assess the biological significance of DE genes near differential boundaries, we examined the biological processes enriched in DE genes near sigDBs compared to processes enriched in other genes (**Methods**). We grouped DE genes into two sets, those near (i.e., within 100kb of) sigDB and those not near sigDB, and tested each set for enrichment of Gene Ontology (GO) biological processes based on FDR-corrected hypergeometric test (**Methods**). In the cardiomyocyte differentiation data, DE genes in both sets showed significant enrichment for generic developmental terms like multicellular organismal development (**Figure 5C**, **Supp Table 8**). However, DE genes near sigDBs tended to be significant for processes specific to cardiac and heart development (e.g. cardiac cell differentiation, heart development and morphogenesis). DE genes near sigDB between H1 and endoderm also showed significant enrichment in developmental terms (e.g. cell morphogenesis involved in differentiation, cellular component organization or biogenesis) compared to those not near sigDB (**Supp Table 9**). For the mouse ESC to CN differentiation, DE genes near sigDB were enriched for neuronal processes when comparing ES vs CN and ES vs NPC (**Supp Table 10**).

Finally, to characterize specific loci with differential 3D organization pattern, we prioritized regions based on the magnitude of change in their boundary scores, then overlapped them with genomic features such as retrotransposons and proximity to DE genes. Human endogenous retrovirus subfamily retrotransposons (HERV-H) in particular have been implicated in chromatin organization (Lawson et al., 2023) as a major determinant of TAD boundaries specific to hESC (i.e., day 0 of cardiomyocyte differentiation) when transcriptionally active (Zhang et al., 2019). We obtained genomic regions with the top 100 most transcriptionally active HERV-H sites and aggregated their TGIF-DB boundary scores at each timepoint. The boundary score at these transcriptionally active HERV-H sites is highest in the hESC (day 0) than in any subsequent timepoints (**Figure 6A**). This is further supported by the presence of a boundary unique to day 0 that disappears in subsequent timepoints at one of the top transcriptionally active HERV-H sites (**Figure 6B**). Among the top-ranked sigDBs based on change in boundary scores, we found sigDB regions in which a boundary is present in the pluripotent state (day 0 hESC state of cardiomyocyte differentiation and H1 cell line, **Figure 6C,D**, respectively), but absent in differentiated state (day 2 mesoderm and definitive endoderm, respectively). These sigDB instances are proximal to the *ESRG* gene, significantly higher in expression in the pluripotent state compared to the subsequent differentiated states in both datasets. *ESRG* is a HERV-H containing long non-coding RNA (lncRNA, Wang et al., 2014); in addition to demarcating domain boundaries in hESCs, this particular site may effect the pluripotency state on knockdown (Wang et al., 2014) and has known roles in developmental and embryonal carcinoma (Wanggou et al., 2012). Among other top-ranked sigDBs in cardiomyocyte differentiation, we found DE genes with known roles in the cardiac development; for example, a particular boundary was found in primitive cardiomyocytes (day 15) but absent in fully differentiated ventricular cardiomyocytes (day 80, **Supp Figure 12A**). This boundary overlaps *MYH6*, highly expressed in day 15 compared to day 80, and is adjacent to *MYH7*, which displays the opposite expression change pattern to *MYH6*. Both these genes are involved in cardiac muscle function: *MYH6* is expressed at high levels in developing atria, while *MYH7* in ventricular chambers of the heart (Ching et al., 2005; Warkman et al., 2012). Recently an enhancer region cluster, located downstream to *MYH7* at chr14:23,876,121-23,878,188, was identified as a switch that can downregulate expression of *MYH7* while upregulating *MYH6* expression upon deletion (Gacita et al., 2021). Further using the same prioritization scheme in mouse neural differentiation data, we found a sigDB close to the *Ncam1* gene, which is differentially expressed between ES and CN (**Supp Figure 12B**); *Ncam1* has known roles in neuron axon guidance and synapse formation (Hata et al., 2018; Shetty et al., 2013). These examples provide further evidence for TGIF-DB’s ability to identify relevant dynamic boundaries that could impact overall cell state identity.

**Figure 6.**
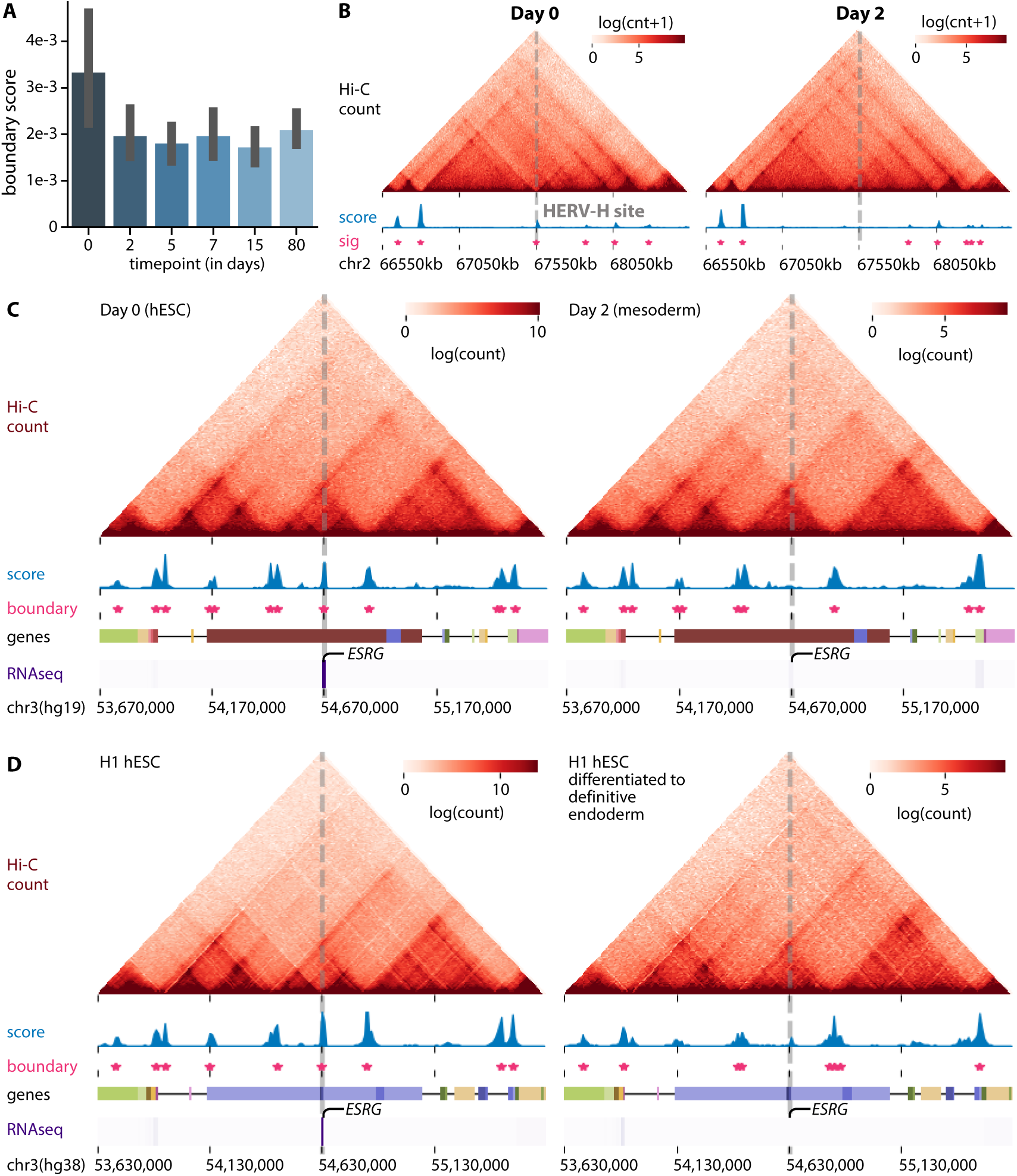
Human pluripotency-specific boundary elements. **(A)** Boundary scores of transcriptionally acive HERV-H retrotransposon sites during each timepoint of cardiomyocyte differentiation. **(B)** The top HERV-H site based on its transcription level in day 0 pluripotent state (within somatic chromosomes) and the overlapping sigDB identified by TGIF-DB. **(C)** A sigDB overlapping ESRG, an HERV-H-containing DE gene, in cardiomyocyte differentiation and in **(D)** H1 differentiated to endoderm.

### Persistent boundaries are enriched for SNPs from diverse disease phenotypes

Single nucleotide polymorphisms (SNPs) identified from genome-wide association studies (GWAS) are frequently found in non-coding regions of the genome and have been implicated in disease phenotypes by affecting the 3D genome organization (Lupiáñez et al., 2015; Orozco et al., 2022). Specifically, such variants could disrupt TAD boundaries and cause promiscuous expression of genes (Lupiáñez et al., 2015; Chakraborty and Ay, 2018). We investigated whether TGIF boundaries from the human cardiomyoctye differentiation data could be used to examine regulatory variants identified for diverse disease phenotypes in GWAS. We considered 17 phenotypic categories from the GWAS catalog and tested the enrichment of SNPs from each category in TGIF boundaries (**Methods**). SNPs across different categories were most enriched in the common set of boundaries across timepoints (i.e., persistent boundaries) than in other timepoint-specific or broader subsets of boundaries, with hematological measurement, cardiovascular disease, and lipid or lipoprotein measurement being the most enriched phenotypic categories (**Figure 7A**). Importantly, SNPs associated with cardiovascular disease (CVD) exhibited the second highest enrichment. The traits that had lower enrichment included neurorological disorders and non-specific categories. We examined 66 persistent boundaries with at least one CVD-associated SNP. One such boundary had the SNP, rs72705895, which is associated with venous thromboembolism (Lindström et al., 2019, **Figure 7B**), and additionally overlaps a CTCF binding site (regulatory feature ENSR00000255184 from Ensembl regulatory build annotations; Zerbino et al., 2015; Cunningham et al., 2022). Another boundary included rs9349379, which is found in the intronic region of *PHACTR1* (**Supp Figure 13**). Both the intronic variant and the gene are associated with coronary artery atherosclerotic disease (Kuveljic et al., 2021; Koitsopoulos and Rabkin, 2021), while the SNP itself is on a predicted enhancer region (Ensembl regulatory build annotation ENSR00001107203), suggesting its putative role in disrupting an intronic enhancer. Genome editing experiments of boundary locations harboring these SNPs combined with Hi-C assays could help examine the role of dysregulated 3D genome organization as a possible mechanism by which regulatory variants impact phenotype.

**Figure 7.**
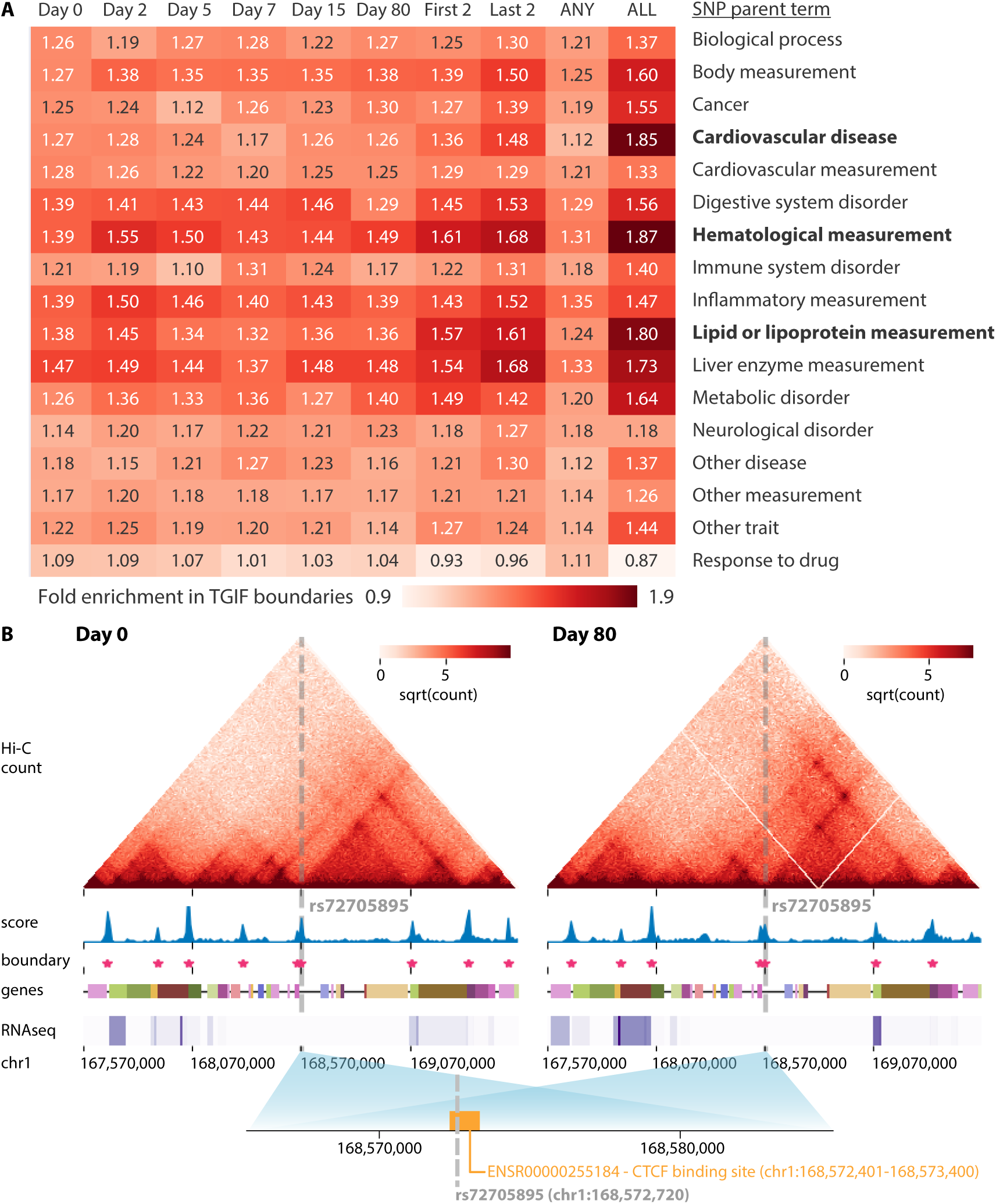
SNP enrichment in persistent boundaries. **(A)** Fold enrichment of SNPs in different subsets of boundary regions, across different categories (SNP parent terms). We measured the enrichment of SNPs in timepoint-specific boundaries (day 0-80) of cardiomyocyte differentiation, in boundaries common to the first 2 or the last 2 timepoints, in union of boundaries (ANY), and in intersection of boundaries across all timepoints (ALL). **(B)** A SNP landing in a boundary persistent across all timepoints (only day 0 and day 80 visualized here) and a CTCF binding site.

## Discussion

Systematic characterization of the dynamics of three dimensional genome organization is important to improve our understanding of how this layer of regulation impacts phenotypic and molecular changes across different biological contexts, such as species, time, and developmental stage. Advances in genomic tools and concerted consortia-level efforts have produced a growing compendia of high-throughput chromosome conformation capture datasets (Dekker et al., 2017, 2023; Reiff et al., 2022). However, systematic analysis of these datasets to quantify the extent of change is a challenge because of the multiple layers at which the 3D genome is organized and the paucity of tools to analyze datasets from a large number of contexts. To address this challenge, we developed Tree-guided Integrated Factorization (TGIF) that combines multitask learning with matrix factorization to examine dynamics of 3D genome organization across multiple structural scales and biological conditions.

TGIF’s design is motivated by a number of considerations: (a) TAD and compartment identification are unsupervised learning problems with no ground truth for real Hi-C datasets. Since Hi-C data can be sparse, identification of such structures and assessing how much they change could be susceptible to statistical, non-biological differences. (b) Several studies from multiple cell types, time points, and species have shown that TAD and compartment is conserved across species (Dixon et al., 2012; Vietri Rudan et al., 2015). TGIF’s hierarchical, multi-task learning framework exploits this prior information to constrain the identification of organizational structures while being sensitive to the extent of relatedness of the datasets by using a tree structure. Thus datasets that are further apart would be constrained less to share similar TAD structure compared to more closely related ones. (c) Finally, TGIF is motivated by a dimensionality reduction (matrix factorization) framework to reduce the noisy, high-dimensional count profile of each genomic locus into a low dimensional space of different ranks. This enables TGIF to be a general framework that identifies TADs, compartments, as well subcompartments and their dynamics. In contrast, existing approaches to identifying differences across multiple Hi-C datasets involve defining structural units of interest independently in each dataset, followed by a post-hoc comparison, or considering pairwise differences. Application of TGIF and existing methods to three mammalian differentiation time course datasets showed that TGIF can accurately recover structural units such as compartments and topologically associated domains (TADs), while being robust to technical differences between datasets such as depth, normalization, and resolution. TGIF also identifies biologically meaningful differences in 3D genome organization that are supported by numerous one-dimensional features such as architectural protein enrichment, histone modification, and differential expression. By allowing users to specify the extent to which the datasets are related, TGIF does not overly impose similarity between datasets. In situations where the datasets are not closely related, we expect TGIF-DB and TGIF-DC to perform similarly to methods that detect TADs (or compartments) per condition and identify differential boundaries and compartments as a post-processing step.

An open question with topological domain changes is how they relate to changes in gene expression (Greenwald et al., 2019; Ghavi-Helm et al., 2019; Cavalheiro et al., 2021; McArthur and Capra, 2021). At the TAD level, fusion or inversion of TADs could result in gene expression change although the extent to which such changes are genome wide or are specific to disease-associated genes is still unclear (Cavalheiro et al., 2021). Evidence suggests that RNA polymerase elongation or the binding of preinitiation complex to the DNA during transcription can give rise to domain structures, providing a direct mechanistic link between transcription and 3D genome organization (van Steensel and Furlong, 2019; Heinz et al., 2018). This relationship can further depend upon the developmental stage or differentiation status of cells (Pollex et al., 2024; Chen et al., 2024). However, this has been debated in other studies, for example, during Drosophila development (Ing-Simmons et al., 2021; Espinola et al., 2021).

Using multi-sample mammalian datasets, we examined the propensity of differentially expressed genes to be close to differential boundaries and compartments. The enrichment of differentially expressed genes near differential boundaries is indicative of the impact of TAD changes to gene expression changes; furthermore, DE genes that were near differential boundaries were more significantly enriched for context-specific processes which could indicate that such changes are associated with fine tuning of gene expression during cellular differentiation. The types of TAD changes we investigated were gain and loss of boundaries between pairs of time points and thus the expansion or contraction of a particular domain. Finally, we observe a similar trend to hold for regions participating in differential compartments, though to a lesser extent that TAD changes. Follow up experiments that perturb boundaries and compartment structures coupled with gene expression measurements would be beneficial for teasing apart causal versus correlational relationships between chromatin organization and gene expression changes.

Regulatory sequence variants can mis-regulate gene expression by disrupting TAD boundaries (Lupiáñez et al., 2015; Chakraborty and Ay, 2018). We used our TAD boundaries to examine the impact of this variation. Interestingly, when considering different types of boundaries based on whether they were timepoint specific, or conserved to different extents, we found the greatest enrichment in boundaries that did not change over time, namely the persistent boundaries. Furthermore, we found several cardiovascular and metabolic disease trait SNPs to be enriched in these boundaries. These persistent boundaries may be specific to the entire cardiac tissue as a whole rather than a specific developmental time or stage. As future work, it would be worth investigating persistent boundaries in other developmental lineages and their propensity to prioritize SNPs for diseases in tissue-specific manner. Additionally, this provides a way to prioritize variants for downstream functional experiments that could be important to identify the mechanisms by which variants disrupt gene regulatory processes.

There are a number of directions in which TGIF could be extended. TGIF expects the relatedness of the datasets to be provided as user input. This information may be available for specific problem settings or can be estimated from the input count matrices in a data-driven way. However, the same hyperparameter is used for all branches of the tree. For situations in which a more granular control is needed between datasets, a direction of future research is to consider the hyper-parameter to vary depending upon the position in the hierarchy. Such information could be informed by auxiliary information such as phylogenetic branch length across species or gene expression similarity across cell types. A second direction of research is to consider auxiliary measurements, including sequence, to inform the inference of the topological units using techniques such as semi-supervised clustering (Bair, 2013; Bondell and Reich, 2008).

Overall, TGIF is a flexible and robust framework to examine changes in genome organization at the compartment and TAD level across a large number of Hi-C datasets. As more datasets across diverse biological contexts become available, methods like TGIF are expected to be increasingly helpful to examine 3D genome organization dynamics and its impact on normal and disease processes.

## Materials and Methods

### Tree-Guided Integrated Factorization (TGIF)

Tree-Guided Integrated Factorization (TGIF) is based on multi-task non-negative matrix factorization (NMF) that can be used to identify low dimensional structure across multiple Hi-C datasets. Below we describe the TGIF framework in detail, which consists of NMF, hierarchical multi-task learning with tree-based regularization, and optimization with block coordinate descent.

NMF is a powerful dimensionality reduction method that can recover the underlying low-dimensional structure from high-dimensional data (Lee and Seung, 2000). It aims to decompose a non-negative matrix, 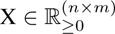, into two lower dimensional non-negative matrices, 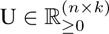 and 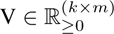, to minimize the following objective: 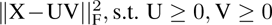, where *||·||*_F_ indicates the Frobenius norm. We refer to the U and V matrices as factors. Here *k ≪ n, m* is the rank of the factors and is an input parameter. As described previously (Lee and Roy, 2021), to apply NMF to Hi-C data, we represent the Hi-C data for each chromosome as a symmetric matrix X = [*x_ij_*] *∈* R^(^*^n×n^*^)^ where *x_ij_* represents the contact count between region *i* and region *j*.

TGIF implements multi-task NMF, where tasks correspond to Hi-C datasets that in turn are from hierarchically related contexts, such as cellular stages, species, timepoints. We note that a hierarchy is a general form for capturing relationships among a set of conditions and can capture both branching and linear relationships. Multi-task NMF has been previously implemented in the multi-view NMF approach (Liu et al., 2013; Baur et al., 2022), where a view and task can be used interchangeably. However, this existing framework assumes that all the tasks are equally related. Formally, in multi-view NMF, given *T* different datasets {X^(1)^ … X^(*T*)^} where each dataset 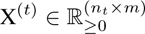, the goal is to find view-specific factors {U^(1)^*, · · ·,* U^(^*^T^* ^)^} and {V^(1)^ … V^(^*^T^* ^)^}, and a consensus factor V*^∗^* that minimize the following objective:

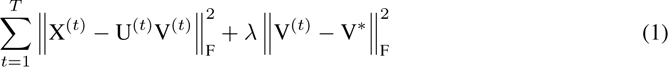

where 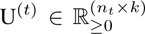 and 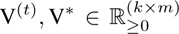. This constrains each of the task-specific factors V^(^*^t^*^)^ to be similar to the consensus factor V*^∗^*. The hyper-parameter *λ* controls the strength of this constraint. The key benefit of such a framework is that the latent representation or structure within each task can borrow from other complementary data, as guided by the consensus factor V*^∗^*.

TGIF generalizes multi-view NMF to allow for integration of datasets that can come from different biological contexts such as time or developmental stage, and therefore may not all be equally related to each other. Accordingly, instead of requiring all the V^(^*^t^*^)^ to be similar to a single V*^∗^*, in TGIF we account for the heterogeneity of the datasets by modeling the tasks to be related by a tree or a hierarchy. This makes TGIF applicable to a wide variety of task collections representing different biological contexts with arbitrary and complex relationships (e.g. Hi-C datasets from different cancer subtypes, cell lineage). In TGIF, the leaves of the tree correspond to the observed dataset while the internal node describe which tasks are most related. The child tasks are then regularized to its immediate parent task.

Formally, TGIF takes as input *t ∈ {*1*, …, T }* tasks, each with input matrix X^(^*^t^*^)^ *∈* R*^n×n^* representing a symmetric count matrix for a Hi-C dataset over *n* genomic loci and a user-specified task hierarchy (**Figure 1A**) and optimizes the following objective:

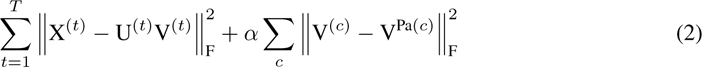

The tree describes parent-child relationships between the tasks. This objective aims to:

1. constrain a task-specific latent factor V^(^*^t^*^)^ in a leaf node of the task hierarchy to be similar to V^Pa(^*^t^*^)^ in its parent node;
2. constrain an internal node’s latent factor V^(^*^b^*^)^ to be similar to its direct child nodes’ *V* ^(^*^c^*^)^ and its parent node’s V*^P^ ^a^*^(^*^b^*^)^;
3. constrain the root node’s latent factor V^(^*^r^*^)^ to be similar to all of its direct child nodes’ *V* ^(^*^c^*^)^s.

The hyper-parameter *α* controls the strength of the constraints such that the higher the *α*, the more the factor V^(^*^c^*^)^ is encouraged to be similar to its parent. If all tasks share a single parent node (the root node), TGIF reduces to multi-view NMF.

Many optimization algorithms exist for learning the latent factors in matrix factorization (Kim et al., 2014). We chose a block coordinate descent (BCD) because it guarantees convergence to a local optimum (Kim et al., 2014). Intuitively, block coordinate descent updates a given block while keeping all other blocks fixed; in TGIF the block is each column or row of U^(^*^·^*^)^s or V^(^*^·^*^)^s. Starting at the leaf nodes (i.e., input tasks), we update each column/row of the factors, then move up the tree to update the parent factors. The specific update rules and their derivations can be found in **Supplementary Methods**.

TGIF’s factors can be used to define compartments as well as fine-scaled topologically associating domains (TADs). TGIF-DC for compartments and TGIF-DB for boundaries and described in detail in subsequent sections.

### TGIF-DB for differential boundary identification

TGIF-DB identifies TAD boundaries in four major steps: (a) multi-task factorization of input Hi-C matrices to obtain low dimensional representations; (b) compute a “boundary” score across different scales, (c) empirical p-value calculation and FDR correction to detect significant boundaries; and (d) determine significant differential boundaries (sigDB) using z-score of pairwise boundary score differences.

#### Multi-task factorization of input Hi-C matrices

TGIF-DB operates on small partially overlapping submatrices along the diagonal of the symmetric intra-chromosomal interaction count matrices (**Supp Figure 1A,B**). This mirrors the approaches taken by existing TAD-calling methods (Lieberman-Aiden et al., 2009; Cresswell and Dozmorov, 2020; Li et al., 2021). By default each submatrix spans 2Mb*×*2Mb with an overlap “step size” of 1Mb between consecutive submatrices. The exact dimension of the submatrix, namely the number of rows and columns, will depend on the resolution of the Hi-C data. For example, for a Hi-C dataset at 10kb resolution, the submatrix will be 200 *×* 200 with an overlapping step size of 100 rows/columns. The minimum size of the submatrices is bound at 100 (and the corresponding step size at 50) to prevent over-fragmentation of the input matrices, especially for lower-resolution input Hi-C matrices. Regions with interaction values missing for more than half of its neighbors in the radius defined by the window size in any of the input matrices are filtered out from the original input intra-chromosomal matrices before any submatrices are formed.

In NMF, usually the rank *k* of the lower dimensional factors is user-specified. However, TGIF does not require this since a single *k* value may not be appropriate across all task-specific input submatrices. Instead TGIF scans a range of *k* values, with *k ∈ {*2*, · · ·,* 8} to recover lower dimensional factors at multiple resolutions and defines boundaries based on a consensus of these factors (as described below). Because the submatrix size is small, it is computationally tractable to scan a range of *k*.

#### Boundary score calculation

After factorization, the next step is to identify genomic regions representing conserved or dynamic TAD boundaries across conditions. We define a boundary as a region whose low-dimensional representation changes significantly compared to its immediate preceding neighbor bin. To this end we define a boundary score 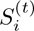 using the output factors for each of the *T* tasks from TGIF. Since **X**^(^*^t^*^)^ is symmetric, either U^(^*^t^*^)^ or V^(^*^t^*^)^ could be used to estimate these boundary scores. Assuming we use U^(^*^t^*^)^, the score 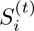 for each region *i* in task *t* is the cosine distance between the low dimensional representation of region *i* and region *i −* 1:

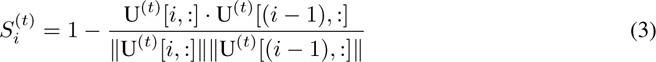

While Euclidean distance could be used to compute the score, we used cosine distance since it normalizes the score regardless of magnitude differences across the tasks (which can arise from Hi-C sequencing depth differences). The final boundary score for region *i* in task *t* is the mean of 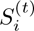 estimated from factors across the range of *k ∈ {*2*, · · ·,* 8}. For regions that are in the overlapping window between two consecutive count submatrices, the final boundary score is averaged from both submatrix factors. This allows us to capture the “boundary-ness” of each region at different structural scales (**Supp Figure 1C**).

#### Empirical p-value calculation and FDR correction

Once the scores are calculated, we estimate a “null” distribution of boundary scores and use it to determine the empirical p-value of boundary scores and find significant boundaries. The null distribution is computed from a randomized background matrix. We first calculate the element-wise mean across *T* input submatrices to yield M, then create randomized background matrix by shuffling the entries of M. The shuffling is done in a distance-stratified manner; that is, we obtain all pairs of genomic regions that are at a distance *d* and permute them independent of the region pairs that are at different distance than *d*. The distance *d* ranges from 0 to the size of each window (e.g. 2Mb), incremented by the size of each bin (e.g. 10kb). We next performed single-task NMF on this shuffled M matrix for the same range of *k* factors and derive boundary scores for all regions in the same way described in **Equation 3**. We treat this set of boundary scores as the samples from the null distribution. We calculate the empirical p-value for each the region *i* in task *t* as the proportion of “null” background scores higher than the given region’s boundary score. Finally, to find significant boundaries and to correct for multiple significant testing, we perform the Benjamini-Hochberg procedure (Benjamini and Hochberg, 1995).

The output of the p-value and FDR estimation step is a binary value for each region *i* and task *t*, indicating whether the region has a significant boundary score (1) or not (0). The significant boundaries identified in this manner may still be susceptible to noisy, low count regions of the genome. Therefore, we additionally filter the boundaries to find “summit-only” version of the significant boundaries, i.e. if there are more than one consecutive significant boundary regions along the linear genome, only the region with the highest significant score is called a boundary.

#### Significantly differential boundary regions in pairwise comparison of conditions

We provide a statistically significant subset of pairwise differential boundary regions (sigDB). For a pair of conditions with input Hi-C matrices, A and B, and for each genomic region *i*, we calculate the absolute difference in boundary scores 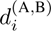 between the two conditions. We estimate a null Gaussian distribution using the absolute difference of boundary scores of genomic regions which do not have significant boundaries in either A and B. We calculate the z-score and corresponding p-value of 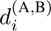 for all regions using this null distribution. After FDR correction, we report the regions with adjusted p-value *<* 0.05 as significantly differential boundary regions.

#### Hyper-parameter selection for TGIF-DB

To determine the setting of *α*, we examined the agreement between boundary assignments from a pair of input matrices for a given *α* value and the similarity of the input matrices themselves (SCC, Yang et al., 2017, **Supp Figure 14A**). We used the cardiomyocyte differentiation data and scanned multiple hyper-parameter values in *α ∈ {*10^2^, 10^4^, 10^6^, 10^8^} across all chromosomes and compared the resulting boundary sets between every pair of timepoints using Jaccard index. In parallel, we measured the similarity of interaction count matrices between every pair of timepoints across all chromosomes using SCC, which is a weighted mean of correlation between interaction counts, stratified by genomic distance. SCC enables unbiased measurement of similarity between Hi-C datasets which are heavily distance-dependent (i.e., closer genomic regions tend to have higher interactions). Finally, we measured the correlation between the Jaccard index and the SCC within each chromosome, across all pairs of timepoints (**Supp Figure 14B,C**). We find a slight, though not significant, improvement with *α* = 10^6^, which we set as default for TGIF-DB.

As BCD is a stochastic algorithm that can reach different local optima depending on the initialization point, we also experimented with multiple random initialization seeds. We used Jaccard index to measure the agreement between pairs of boundary sets from two different seeds, with *α* fixed at the default value 10^6^. We found that the resulting boundary sets from different initialization are relatively consistent with pairwise Jaccard index 0.76-0.77 (**Supp Figure 14D**). We also estimated the memory usage and run time trend of TGIF-DB on 10kb input matrices from the three different timecourse datasets (**Supp Figure 15A,B**). TGIF-DB’s submatrix factorization framework with fixed set of *k* makes it scale linearly with the size of the input matrices.

### TGIF-DC for differential compartment and subcompartment identification

#### Identification of compartments with TGIF-DC

In order to identify compartments, we apply TGIF to a 100kb resolution intrachromosomal Hi-C matrix that is first converted into an observed-over-expected (O/E) count correlation matrix as described previously (Rao et al., 2014, **Datasets and preprocessing**). To obtain the O/E matrix, we first remove rows and columns corresponding to regions with zero interactions, then distance-normalize every entry of the input matrices 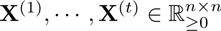 :

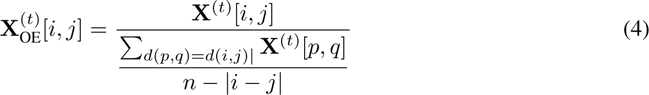

where the denominator is the expected count for pairs of loci *i* and *j* at distance genomic distance *d*(*i, j*). Next we get the correlation matrix as:

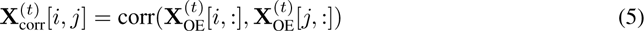

where corr corresponds to the Pearson’s correlation.

We upshift the correlation matrix by 1 so that all values are non-negative. To identify compartments, we apply TGIF with the input tree structure to these matrices with rank *k* = 2. After factorization, we infer each region *i*’s cluster assignment, 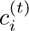, for each task *t*, such that 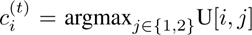. We refer to these clusters as compartments. To identify subcompartments and differential subcompartment regions, a higher *k* value, e.g. 5, can be used and TGIF-DC will generate more granular cluster assignments, e.g. 5 clusters of regions instead of 2 clusters. Each of these clusters corresponds to a subcompartment.

#### Detecting differential compartments with TGIF-DC

While the cluster assignment switch is a straightforward way of identifying differential compartment regions, we further provide a statistically significant subset of pairwise differential compartment regions. We utilize the lower-dimensional representation of each genomic region from the factors in this step. For a pair of conditions or timepoints being compared, A and B, we calculate the cosine distance 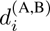 between U^(^*^A^*^)^[*i,* :] and U^(^*^B^*^)^[*i,* :] for each genomic region *i*. Using the cosine distance of regions that do not change their cluster assignment between the conditions (i.e., static regions), we estimate the mean and standard deviation of a Gaussian null distribution. The null distribution is used to calculate the z-score and p-value for the remaining (dynamic/differential) regions. Statistically significant differential regions are those with an FDR *<* 0.05. Significantly differential subcompartment regions are identified in the same way as the differential compartment regions.

#### Hyper-parameter selection for TGIF-DC

To determine the best setting of *α*, we used a similar approach as for TGIF-DB, measuring the agreement (correlation) between the input matrix similarity and the agreement of TGIF-DC compartment assignments for the same inputs. Specifically, we measured the similarity of observed-over-expected (O/E) count matrices between every pair of timepoints across all chromosomes by flattening them into a vector and measuring correlation. Note, O/E counts are already normalized by distance. In parallel, we measured the similarity of the output clusters (compartments) with Rand index (**Supp Figure 16A**). We used the mouse neural differentiation data to study this parameter, scanning *α ∈ {*10^2^, 10^4^, 10^6^, 10^8^} across chromosomes (**Supp Figure 16B**). TGIF-DC consistently yields cluster assignments that are well correlated (*∼*0.9) with the input matrix similarity across wide a range of *α* values (**Supp Figure 16C**). Our results are from *α* = 10^4^, which is the default for TGIF-DC.

Similar to TGIF-DB, we also examined the stability of compartment assignments with multiple random initialization seeds with *α* = 10^4^ using Rand Index between pairs of cluster assignments from two different seeds. At *k* = 2, the compartment setting of TGIF-DC, the Rand Index ranged from 0.99-1 for all pairs of random initializations (**Supp Figure 16D**). At *k* = 5, the subcompartment setting, Rand Index of resulting subcompartments was 0.7-0.8 (**Supp Figure 16E**), showing that TGIF-DC yields a stable set of cluster assignments regardless of random initialization. Finally, we report the memory usage and run time trend of TGIF-DC on 100kb input matrices from the three different timecourse datasets (**Supp Figure 17A,B**). TGIF-DC can analyze 6 datasets needing no more 0.5GB of memory and 400 seconds of run time.

#### Post-hoc annotation of TGIF-DC clusters into A and B compartments

TGIF-DC by default uses *k*=2 and segments the given chromosome into 2 clusters of regions. In our analysis, we use GC content and chromatin accessibility to annotate each cluster as A or B compartment. Note that identification of significant differential compartmental region (sigDC) does *not* depend on this post-hoc annotation step, as sigDC identification only looks at cluster assignment changes and absolute change in latent features of each genomic region.

For the H1 to endoderm differentiation dataset we used the available compendium ATACseq data from 4D Nucleome (Reiff et al., 2022; Dekker et al., 2023, **Supp Table 1**) as a measurement of chromatin accessibility. For each chromosome, we measure the mean ATACseq reads per 100kb bin from H1 within each of the 2 clusters. The cluster with higher mean ATACseq signal is assigned to compartment A and the other to compartment B for both H1 and endoderm (this is possible due to the cluster correspondence across timepoints in TGIF-DC; **Supp Figure 18A,B,C**). To validate the compartment annotation, we also measured the GC percentage within each 100kb bin for each compartment, and found that regions assigned to A compartment have higher GC content than those in B compartment, as expected (**Supp Figure 18D**).

We proceeded similarly with cardiomyocyte differentiation data, except we used DNase-seq data for chromatin accessibility from day 0 (H9 cell line, hESC state, **Supp Table 3**). We measure the mean DNase-seq reads per 100kb bin for both clusters in each chromosome. The cluster with higher mean DNase-seq signal is labeled compartment A and the other one is labeled compartment B across all timepoints (**Supp Figure 19A,B**). To validate the compartment annotation, we also measured GC percentage and found that regions assigned to the A compartment has higher GC content (**Supp Figure 19C**). Since H3K27ac was available for this dataset, we also compared the mean H3K27ac reads within each 100kb bin of each compartment and found higher H3K27ac signal in the A compartment compared to B (**Supp Figure 19D**). This is consistent with the definitions of A and B compartments (Nichols and Corces, 2021; Bouwman et al., 2023) and provides further support for TGIF-DC’s compartmentalization.

For mouse neural differentiation data, we only used GC content for compartment annotation. We performed this annotation for the ES timepoint and transferred it to the other timepoints. Briefly, for each chromosome, we measure the mean GC% per 100kb bin in each compartment. The compartment with higher mean GC content is called A and the other one B across all timepoints (**Supp Figure 7**).

### Estimating tree structure from input Hi-C matrices for unknown inter-dataset relationships

Typically, prior information about the relatedness of the biological contexts from which the Hi-C data matrix originate is available. However, if this information is not available (e.g. integrating Hi-C datasets from different laboratories or single-cell Hi-C datasets), a tree can be estimated using pairwise similarity of the input Hi-C matrices, converting to distance followed by hierarchical clustering. We suggest the use of a distance-stratified similarity measure, such as the stratum-adjusted correlation coefficient (SCC, Yang et al., 2017, **Supp Figure 2A**), that we have also used for our hyper-parameter analysis. Once SCC is calculated for each pair of input matrices, it is converted to a distance by subtracting from 1 (**Supp Figure 2B**) which in turn is used as input to hierarchical clustering with average linkage. We tested this approach for the mouse neural differentiation dataset and found that the output tree of hierarchical clustering is similar to the known biological relatedness of this dataset (**Supp Figure 2C**, Bonev et al., 2017) and is identical to the tree we used as input to TGIF for our experiments. The current implementation of TGIF offers this functionality as a pre-processing script (See Section on **Data access and code Availability**).

### Datasets and preprocessing

We applied TGIF to three Hi-C timecourse datasets: H1 hESC differentiated to endoderm (Reiff et al., 2022; Dekker et al., 2023), mouse neural differentiation data from Bonev et al., 2017, and human cardiomyocyte differentiation data from Zhang et al., 2019 (**Supp Figure 3**). The Hi-C interaction count matrices for pluripotent H1 hESC cell line and for endoderm differentiated from H1 was downloaded from 4D Nucleome consortium (**Supp Table 1**). 100kb intra-chromosomal count matrices were used for comparison of compartment-calling methods. Additionally, ATACseq data for H1 and endoderm was also downloaded and used to measure the accessibility of each 100kb region (i.e. mean ATACseq reads per base in the region) in the comparison of compartment-calling methods. 10kb VCSQRT-normalized intra-chromosomal count matrices was used in the analysis of differential boundary and expression analysis.

The mouse neural differentiation data (Bonev et al., 2017) included 3 timepoints during mouse neural differentiation: embryonic stem cell (ES) stage, neural progenitor stage (NPC), and the differentiated cortical neuron (CN) stage. For TGIF-DB, we used intra-chromosomal count matrices at a resolution of 10kb resolution with vanilla-coverage square-root (VCSQRT) normalization as input. When benchmarking boundary-calling methods, we also used 25kb and 50kb VC-SQRT-normalized data, as well as 25kb ICE-normalized data. For TGIF-DC, intra-chromosomal interaction count matrices at 100kb resolution without normalization was used as input since TGIF-DC computes the O/E correlation matrices. In addition to the Hi-C measurements, this dataset also included ChIPseq data for 6 histone modification marks (H3K27ac, H3K28me3, H3K36me3, H3K4me1, H3K4me3,H3K9me3), for both CN and NPC (**Supp Table 2**). This data was used to characterize and validate the chromatin structure inferred by TGIF-DC. For each 100kb bin, the ChIPseq signal in reads-per-million was summed up within each bin, and the signal in each bin was divided by the total number of reads to first normalize by read depth. Subsequently, signals in NPC and CN were quantile normalized to each other to enable log fold change comparison across the two timepoints. The log fold change for each of the 6 marks in each bin was calculated by log transforming (log(x+1)) the normalized signal in NPC divided by the signal in CN. The set of log fold change signals were used as input to a k-means clustering, with *k* = 5 clusters applied to sigDC regions.

The cardiomyocyte differentiation dataset from Zhang et al., 2019 (**Supp Table 3**) measured Hi-C counts at 6 different timepoints (day 0, 2, 5, 7, 15, 80) from the human embryonic stem cell stage (hESC, day 0) to the ventricular cardiomyocyte stage (day 80). Two replicates from each timepoint were first merged and intra-chromosomal interaction count matrices were generated at 10kb resolution with ICE normalization using the Juicer tool (Durand et al., 2016). The 10kb resolution matrix was provided as input to TGIF-DB. The merged dataset was also used to generate 100kb resolution intra-chromosomal count matrices for input to TGIF-DC. **Benchmarking with downsampled data to assess robustness to depth** section contains details on how GM12878 cell line data was processed and used.

### Benchmarking methods for identifying differential domain boundaries

#### Description of existing methods

TGIF-DB was benchmarked against four other methods for identifying differential TAD boundaries: GRINCH (Lee and Roy, 2021), SpectralTAD (Cresswell et al., 2020), TADCompare (Cresswell and Dozmorov, 2020), and TopDom (Shin et al., 2016). GRiNCH, SpectralTAD, and TopDom are single-task TAD identification methods accepting a single input matrix individually followed by pairwise comparison of identified boundaries.

GRiNCH is a TAD identification method that also utilizes a variation of NMF. It applies graph-regularized NMF to an input intra-chromosomal matrix as a whole and uses the output factor to cluster the genomic regions; each cluster represents a TAD. The boundary of such clusters were used as TAD boundaries in our analysis. For pairwise differential analysis, boundaries found in both input matrices were considered shared boundaries, while those found in one and not the other were considered differential boundaries. Version 1.0.0 was used with default values for all optional hyperparameters.

SpectralTAD treats an input Hi-C matrix as a graph of interacting genomic regions, and applies eigen decomposition to its graph Laplacian. The eigenvectors are used as latent features of each genomic region to cluster them, with each cluster representing a TAD. Similar to GRiNCH, for pairwise differential analysis, boundaries found in both input matrices were considered shared boundaries; those found in one and not the other differential boundaries. Version 1.16.1 was used in our analysis.

TopDom is a TAD identification method which was shown to be robust to noise and yielded TADs enriched in structural proteins such as CTCF (Dali and Blanchette, 2017; Lee and Roy, 2021). It generates a score for each bin along the chromosome, where the score is the mean interaction count between the given bin and a set of upstream and downstream neighbors (neighborhood size is a user-specified parameter). Putative TAD boundaries are picked from a set of bins whose score forms a local minimum; false positive boundaries are filtered out with a significance test. Differential boundaries are identified in a pairwise manner similar to GRiNCH and SpectralTAD. Version 0.10.1 was used in our analysis, with the window size hyper-parameter set to the recommended value of 5 (Shin et al., 2016).

TADCompare is a differential TAD identification method that can take as input a pair of Hi-C matrices as well as a time series of Hi-C matrices. It treats each Hi-C matrix as a graph where each genomic region is a node and the pairwise interaction is a weighted edge with the weight corresponding to the count. It performs eigen decomposition on the Graph Laplacian of each Hi-C matrix. Boundary scores are calculated between a pair of regions by computing the Euclidean distance between their corresponding rows in the eigenvectors. Differential boundary scores are calculated by taking the difference in the boundary scores for a pair of conditions and converting it into a z-score and using a threshold of 2 to define a differential boundary. Although TADCompare accepts time-series data differential boundaries are computed for only pairs of matrices at a time. Version 1.2.0 was used in our benchmarking analysis.

In addition to the above mentioned tools, we initially considered the TADsplimer (Wang et al., 2020), and HiCExplorer (Wolff et al., 2020) methods. TADsplimer was specifically developed to detect differential TAD identification; however, its implementation has an unresolved issue that fails to return any output or differential boundaries and was excluded from further analysis. HiCExplorer allows TAD finding followed by differential TAD analysis using its hicDifferentialTAD tool. However, hicDifferentialTAD expects the same TAD for different conditions and outputs TADs with significantly different interaction counts rather than detecting boundary changes. Due to these reasons, they were excluded from subsequent benchmarking.

#### Measuring CTCF enrichment in boundaries

To evaluate the boundaries identified by various TAD-calling methods, we measured CTCF peak enrichment in those boundaries on the cardiomyocyte differentiation dataset which had CTCF ChIPseq data and as such represented a dataset with the largest number of time points and different types of assays (**Supp Table 3**). Using macs2 (Zhang et al., 2008), we first called peaks on CTCF ChIPseq data from each of the 6 timepoints (day 0, 2, 5, 7, 15, 80) of the cardiomyocyte differentiation time course. Replicates from each timepoint were collapsed by intersecting overlapping peaks with Bedtools (Quinlan and Hall, 2010). Each peak was then assigned to a 10kb uniform bin again using Bedtools. TAD-calling methods GRiNCH, SpectralTAD, and TopDom were applied to 10kb Hi-C matrices from each of the 6 timepoints to yield 6 timepoint-specific sets of boundary bins. TGIF-DB was applied to Hi-C matrices from all 6 timepoints using the tree structure as in **Supp Figure 3**, and significant boundaries from each timepoint were used for enrichment analysis. TADCompare was applied to each pair of consecutive timepoints: day 0 vs 2, 2 vs 5, 5 vs 7, 7 vs 15, 15 vs 80. As TADCompare outputs both non-differential and differential boundaries for every pairwise comparison, we define a boundary set specific to a timepoint as follows: (1) for day 0, union of differential boundaries in day 0 and non-differential boundaries between day 0 and 2; (2) for day 80, union of differential boundaries in day 80 and non-differential boundaries between day 15 and 80; (3) for all intermediate timepoints *t*, union of differential boundaries in *t*, non-differential boundaries between day *t* and *t*_previous_, and non-differential boundaries between day *t* and *t*_following_.

The CTCF peak fold enrichment ratio for a given timepoint was calculated as 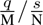, where *q* is the number of boundaries with at least one CTCF peak, *M* is the number of boundary regions, *s* is the number of regions with at least one CTCF peak, and *N* is the total number of genomic regions. We also used these estimates to compute a p-value using a hypergeometric test; however, the fold enrichment was more informative for comparing methods.

#### Benchmarking with downsampled data to assess robustness to depth

We downloaded the high-depth Hi-C dataset of GM12878 cell line (Rao et al., 2014) with 4.01 billion total reads from the 4D Nucleome data portal (Reiff et al., 2022; Dekker et al., 2017, **Supp Table 4**). We then subsampled 5, 10, 25, 50% of the reads, generated 10kb-resolution intra-chromosomal Hi-C matrices using Juicer (Durand et al., 2016), and ICE-normalized the intra-chromosomal interaction matrices from each downsampled dataset (see scripts on https://doi.org/10.5281/zenodo.13323899 for details). We used Jaccard index to measure agreement of boundaries identified by different methods despite depth difference. The Jaccard index is calculated by dividing the number of boundaries found at both depths by the number of boundaries identified in either depths for the GM12878 dataset. The higher the Jaccard Index, the fewer the false-positive differences.

Three TAD-calling methods, GRiNCH, SpectralTAD, and TopDom were applied individually to 5 datasets: original high-depth GM12878 data and four low-depth GM12878 data downsampled to 5, 10, 25, 50% depths, respectively. Jaccard index was calculated between the boundaries identified from the original dataset and those from each of the downsampled datasets. TADCompare and TGIF-DB were applied to 4 pairs of datasets, each pair including the original high-depth GM12878 dataset and the downsampled low-depth dataset (e.g., GM12878 data downsampled to 50% depth, **Supp Figure 3**). For TADCompare, the Jaccard index was calculated for each pair of datasets as the ratio of number of non-differential boundaries and the number of differential and non-differential boundaries. Similarly, for TGIF-DB, the Jaccard index was calculated as the ratio of the number of non-significantly differential boundary regions divided by the size of the union of boundary regions from original-depth and subsampled dataset.

#### Measuring stability of boundary sets across multiple resolutions of input data

We used the mouse neural differentiation dataset (Bonev et al., 2017) to assess the stability of boundary sets identified by different TAD boundary identification methods at different resolutions, namely, 10kb, 25kb and 50kb, since this dataset was readily available at these resolutions. We focused our comparisons only for the mouse embryonic stem cell (mESC) time point. The single-task boundary calling methods (GRiNCH, SpectralTAD, TopDom) were applied individually to 10kb, 25kb, and 50kb intra-chromosomal matrices from mESC. TADCompare was applied to a pair of timepoints including mESC, both at the same resolution: mESC vs neural progenitors (NPC), and mESC vs cortical neurons (CN). In order to find mESC boundaries from the outputs of the pairwise TADCompare comparisons at each resolution, we took the union of non-differential boundaries and differential boundaries enriched in mESC. We applied TGIF-DB to a tree with all three timepoints from mouse neural differentiation dataset at a specific resolution, and took the significant boundaries from mESC (**Supp Figure 3**). This was repeated for each resolution. To allow for comparison of boundaries from different resolutions, we project the higher resolution bins to the coarsest resolution, namely 50kb. For instance, in the 25kb vs 50kb comparison, each 50kb bin is composed of two 25kb bins and is considered to have a boundary if either of the 25kb bins had a boundary. Similarly, for the 10kb vs 50kb comparison, any of the 5 comprising 10kb bins would be used to define a boundary in the 50kb bin spanning them. In the 10kb vs 25kb comparison, if any of the 10kb bins or the 25kb bins have a boundary in the shared 50kb bin, we define the 50kb bin as a boundary. We then measure Jaccard index of boundaries at this lowest resolution.

### Comparison of TGIF-DC to existing compartment-calling methods

We compared TGIF-DC to two established methods for calling compartments, i.e. principal component analysis (PCA) based method (Lieberman-Aiden et al., 2009) and Cscore (version 1.1, Zheng and Zheng, 2018), as well as a method designed specifically for differential compartment analysis, dcHiC (version 2.1, Chakraborty et al., 2022). We applied all four methods to 100kb intra-chromosomal count matrices from H1 hESC cell line. TGIF-DC and dcHiC were additionally applied to 100kb intra-chromosomal count matrices from H1 differentiated to endoderm. Both datasets were downloaded from 4D Nucleome consortium (Reiff et al., 2022; Dekker et al., 2023).

The PCA-based method applies PCA to the observed over expected count (O/E) correlation matrices. The first principal component (PC1) is used to assign genomic regions to two compartments; values equal to or above zero in PC1 are assigned to one compartment and those below zero are assigned to the other compartment. The actual annotation of each compartment as the active “A” or repressed “B” compartments is done by correlating the compartment structure to one-dimensional regulatory signals such as ATACseq assays or histone modifications. Cscore outputs a score that specifies the compartment of a region by modeling the interaction counts and genomic distance with a probabilistic model. Regions with Cscore values above or equal to zero were clustered into one compartment and those below another compartment.

Finally, dcHiC is a framework that can identify differential compartment regions. dcHiC performs fast PCA on distance-normalized correlation matrices, quantile-normalizes the PC values so they can be compared across multiple conditions, and identifies a set of genomic regions whose compartment scores (normalized PC values) are significantly different in any of the conditions using a chi-square test (Chakraborty et al., 2022). As the current version of dcHiC requires at least 2 replicates per condition or timepoint, we provide dcHiC with interaction counts for two replicates per condition. For the other methods we provide a replicate-merged count matrix per condition available from the 4D Nucleome consortium.

To compare the compartment results across the different methods, we measured the Rand index between compartment assignments to each genomic region. To measure the quality of the compartments, we used three well-known cluster quality metrics: Silhouette Index, Calinski-Harabasz Score, and Davies-Bouldin Index, measured on the observed-over-expected (O/E) matrices for each chromosome, as well as the accessibility signal for each 100kb genomic region. The accessibility signal was defined as the mean ATACseq reads per basepair.

Finally, to compare dcHiC and TGIF-DC for significantly differential compartments between H1 and endoderm, we calculated the log ratio of the accessibility signal and gene expression (from RNAseq, in TPM) in H1 over that of endoderm for each significantly differential region.

### Assessing differential gene expression near or within significantly differential boundaries and compartments

We used RSEM (Li and Dewey, 2011) on the raw RNA-seq data from the cardiomyocyte differentiation and the mouse neural differentiation time course to obtain expected counts for each replicate at each timepoint. We also downloaded the RNAseq data for H1 hESC cell line and endoderm differentiated from H1 from 4D Nucleome (Reiff et al., 2022; Dekker et al., 2023). We used these values as input to DESeq2 (Love et al., 2014) to identify differentially expressed (DE) genes for every pair of timepoints in each dataset (e.g., H1 vs endoderm; mESC vs NPC; day 0 vs. day 2 in cardiomyocyte differentiation). DE genes were defined by using a threshold of adjusted p-value *<*0.05.

For every pair of timepoints, we tested the enrichment of these DE genes within regions of interest (**Figure 5A**): (A) regions near (i.e., within 100kb) significantly differential boundaries (sigDB), (B) regions within a TAD with at least one sigDB, and (C) regions within significantly differential compartmental regions (sigDC). For (B), we define all regions bounded within a pair of shared boundaries and containing at least on sigDB within those bounds as belonging to a “TAD with at least one sigDB”.

The fold enrichment of DE genes in these regions was computed as 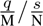, where *N* is number of all regions, *s* = *|*set of regions with at least one DE gene*|*, *M* = *|*a subset regions of interest as defined above, e.g., regions near sigDB*|*, *q* = *|*regions of interest with at least one DE gene, e.g., regions near sigDB with a DE gene*|*. We also performed gene-centric fold enrichment calculations: 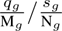, where *N_g_* is total number of genes with expression, *s_g_* = *|*DE genes*|*, *M_g_* = *|*genes overlapping with a regions of interest, e.g., region near sigDB*|*, *q_g_* = *|*DE genes overlapping with a regions of interest*|*. Hypergeometric test was additionally performed to calculate the significance of this fold enrichment value for each pair of timepoints.

Gene ontology (GO) term enrichment analysis was performed for two different subsets of genes based on their DE status and whether they were close to (within 100kb of) sigDB: (1) DE genes not close to a sigDB, (2) DE genes close to a sigDB. The significance of enrichment was determined with an FDR-corrected hypergeometric test p-value *<*0.05. To select candidate differential boundaries for visualization, we ranked a sigDB based on two criteria: (1) adjusted p-value of the change in TGIF-DB boundary score, and (2) the significance of the nearby differential expression measured by the nearest DE gene’s adjusted p-value. We converted these values into ranks and used the mean rank of a boundary to select top 10 regions with promising differential boundaries.

### SNP enrichment within TGIF boundaries from cardiomyocyte differentiation data

We downloaded SNPs in the GWAS catalog (Sollis et al., 2023) and mapped each SNP’s associated trait to its parent phenotype, based on Experimental Factor Ontology (EFO). We refer to these parent terms as SNP categories in our analysis. In total we had 17 such categories (e.g. cardiovascular disease) for which we tested enrichment of SNPs in TGIF-DB boundaries. For each category, we calculated the fold enrichment of associated SNPs in different subsets of TGIF-DB boundaries across different timepoints:

boundaries found in a specific timepoint, boundaries found in the first two or the last two timepoints, boundaries found across all timepoints (ALL, **Figure 7A**), boundaries found in any of the timepoints (ANY, **Figure 7A**). We used the following formula to calculate fold enrichment: 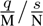. Here, *q* is the number of boundaries of a particular type (e.g. ANY) with at least one SNP of interest, *M* is the number of boundaries of a particular type (e.g. ANY), *s* is the number of regions containing at least one SNP, and *N* is the total number of genomic regions.

## Data access and code availability

TGIF-DC and TGIF-DB are available at https://github.com/Roy-lab/tgif. All scripts used for evaluation, analysis, and visualization have been deposited at https://doi.org/10.5281/ `zenodo.13323899`. See **Supp Table 1-4** for the detailed list of accession numbers for all data used in our analysis.

## Competing Interests

Authors have no competing interests.

## Acknowledgements

This work is supported by the National Institutes of Health (NIH) through the grant NIH NHGRI R01HG010045-01 and by the Computation and Informatics in Biology and Medicine (CIBM) training program (NLM 5T15LM007359). We thank the Center for High Throughput Computing at University of Wisconsin - Madison for computational resources. We also thank Yanxiao Zhang and Bing Ren for providing the list of HERV-H retrotransposon site coordinates and their expression levels.

## Author contributions

Lee and Roy conceptualized the overall framework and algorithm. Lee implemented the algorithm, designed and performed experiments, and wrote the manuscript. Roy designed the experiments and wrote the manuscript.

## Notes

https://github.com/Roy-lab/tgif

https://zenodo.org/doi/10.5281/zenodo.13323898

### Competing Interest Statement

The authors have declared no competing interest.

### Summary of Updates

The manuscript has been updated with expanded benchmarking and methodological extensions to TGIF: 1. extensive benchmarking of both TGIF-DB and TGIF-DC with more methods and metrics in real Hi-C data; 2. extension of TGIF-DB and TGIF-DC to identify significantly differential regions; 3. additional feature in TGIF-DC to call more granular compartments, i.e., subcompartments; 4. an optional pre-processing step to infer a task hierarchy based on input matrix similarity. In total, our revisions have resulted in major additions and updates to 5 main figures, addition of 15 new supplementary figures and 9 new supplementary tables, as well as corresponding updates and clarifications in the main text for results, methods, and discussion points.

