## Supplementary Materials for "Examining dynamics of three-dimensional genome organization with multi-task matrix factorization"

### 1 Supplementary Figures

#### 1.1 Supp Figure 1

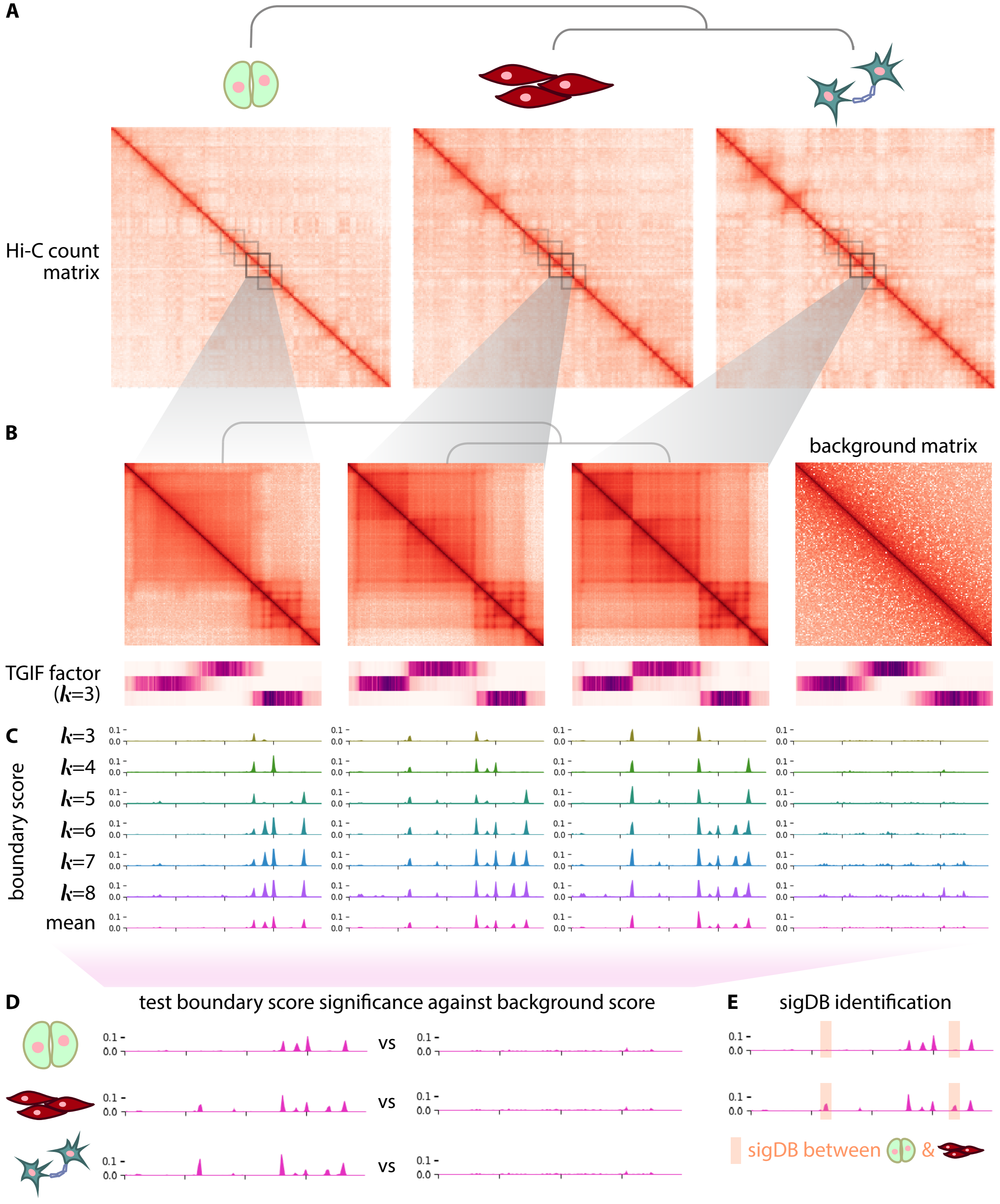

**Supp Figure 1.** Overview of TGIF-DB. **(A)** TGIF-DB takes as input a user-specified tree structure that encodes the relationship among the input Hi-C datasets. Shown is an example of a tree encoding a cell lineage across 3 cell types. **(B)** Multitask matrix factorization is performed for submatrices along the diagonal of the input intra-chromosomal Hi-C matrices. **(C)** Within each submatrix, a range of hyper-parameter $k$ values is used to generate boundary scores for TADs at different scales. For each genomic region, the mean boundary score is taken across the range of $k$s. **(D)** The mean boundary scores are compared to a "null-distribution" boundary scores generated from a randomly shuffled matrix in order to calculate an empirical p-value for each genomic region and to identify significant boundaries. **(E)** Significantly differential boundaries (sigDB) are identified for every pair of input matrices (only 1 pair shown here as example).

#### 1.2 Supp Figure 2

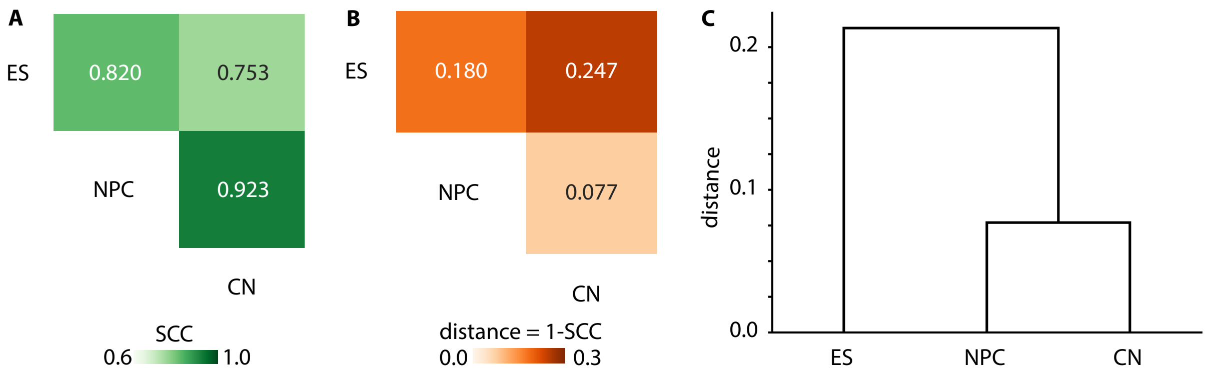

**Supp Figure 2.** Inferring a tree structure between input conditions based on the similarity of the input matrices. Here we use mouse chr19 intra-chromosomal matrices (25kb resolution) from neural differentiation data with 3 timepoints (ES, NPC, CN) as an example. **(A)** Similarity between input matrices for each pair of timepoints, measured by stratum-adjusted correlation coefficient (SCC). **(B)** Similarity converted to distance = 1-SCC. **(C)** Dendrogram of hierarchy constructed using the distance and average linkage.

#### 1.3 Supp Figure 3

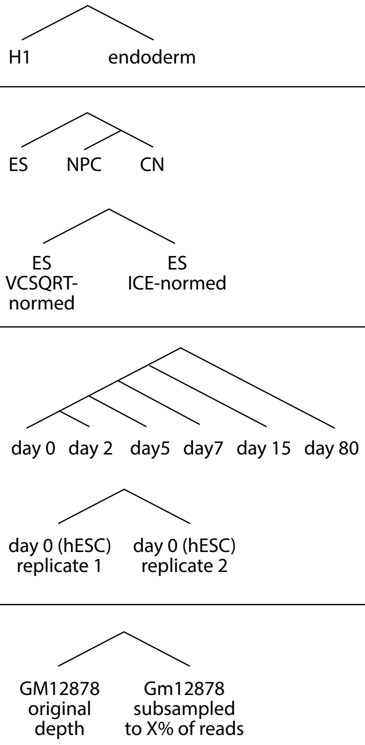

| dataset | inputs/timepoints | tree structure | used in analysis |
|---|---|---|---|
| H1-endoderm (Reiff et al., 2022, Dekker et al., 2023) | H1 (hESC), definitive endoderm differentiated from H1 | H1, endoderm | benchmarking: compartment -calling & differential compar-tment methods, DE analysis |
| mouse neural differentiation (Bonev et al., 2017) | ES (mESC), NPC (neural progentors), CN (cortical neurons) | ES, NPC, CN | benchmarking: TAD stability to resolution, compartment histone mark charaterization, DE analysis |
| | | ES VCSQRT-normed, ES ICE-normed | benchmarking: TAD stability to normalization method |
| cardiomyocyte differentiation (Zhang et al., 2019) | day 0 (hESC), day 2 (mesoderm), day 5 (cardiac mesoderm), day 7 (cardiac progenitors), day 15 (primitive cardiomyocytes), day 80 (ventricular cardiomyocytes) | day 0, day 2, day5, day7, day 15, day 80 | benchmarking: CTCF enrich-ment in TAD bondaries, DE analysis, SNP analysis |
| | | day 0 (hESC) replicate 1, day 0 (hESC) replicate 2 | benchmarking: TAD stability across biological replicates |
| GM12878 cell line (Rao et al.,2014) | GM12878 reads subsampled to 5, 10, 25, 50% of original depth | GM12878 original depth, Gm12878 subsampled to X% of reads | benchmarking: TAD stability to read depth |

**Supp Figure 3.** Overview of datasets used in benchmarking and analysis.

#### 1.4 Supp Figure 4

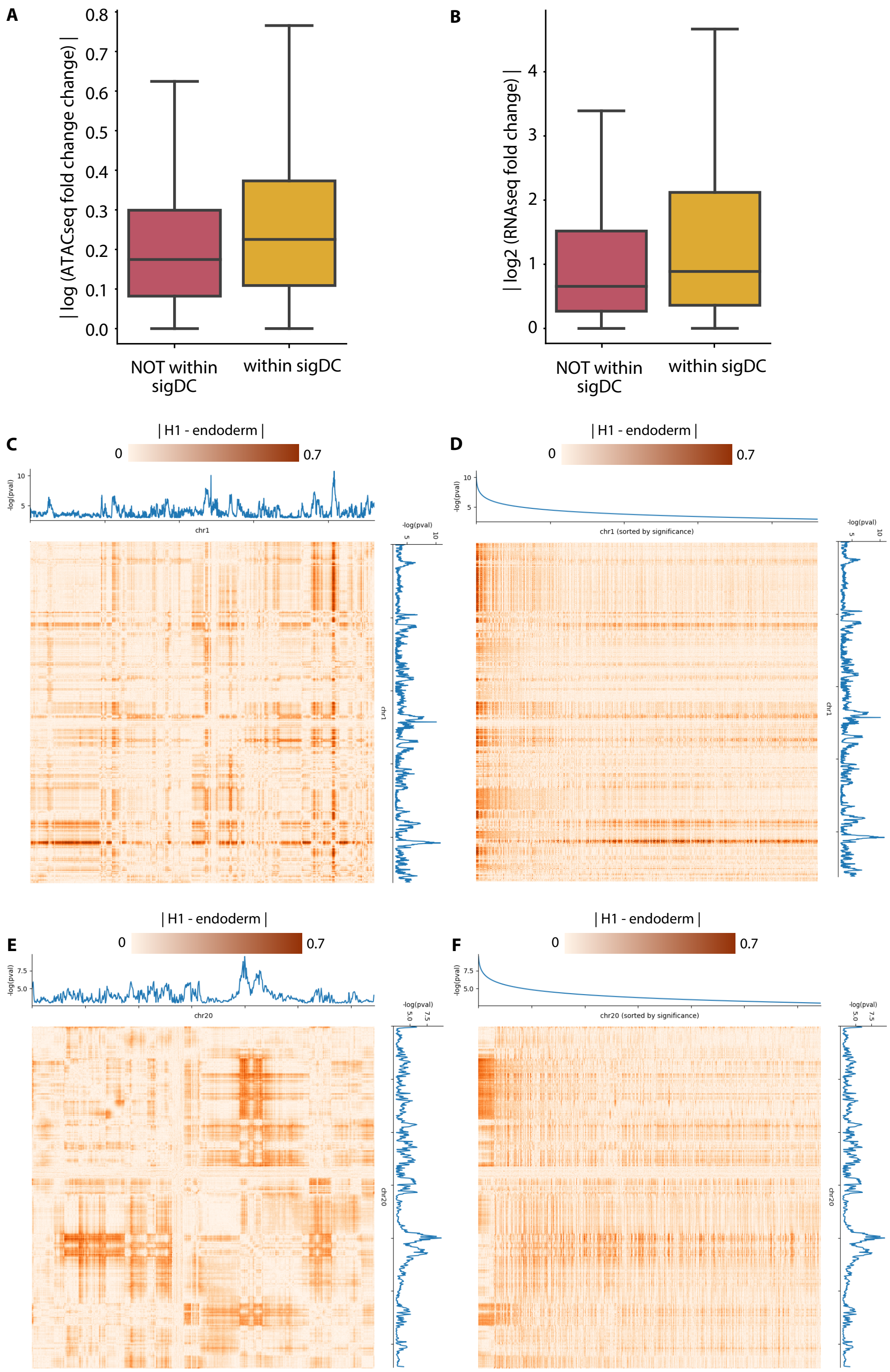

**Supp Figure 4.** Characterizing sigDC in H1-endoderm differentiation. **(A)** Changes in accessibility within significantly differential compartment regions (sigDC). **(B)** Changes in gene expression level within sigDC. **(C)** Visualization of the difference in the input matrices (heatmap) and the significance of differences estimated with TGIF-DC (lineplot). Each row and column of the heatmap is a 100kb genomic region of chr1 and each entry in the heatmap = $corr(O/E)_{H1}$ - $corr(O/E)_{endoderm}$. The lineplot shows -log(adjusted p-value) from TGIF-DC used for detecting significantly differential compartment regions between H1 and endoderm. **(D)** Heatmap shows the same information as in **(C)**, but with columns and rows sorted in decreasing significance, i.e., TGIF-DC's -log(adjusted p-value). The sorting of regions by p-value highlights greater differences in count for regions with higher negative log p-values (high significance). **(E)** Same visualization as **(C)**, but for chr20. **(F)** Same visualization as **(D)**, but for chr20.

##### 1.5 Supp Figure 5

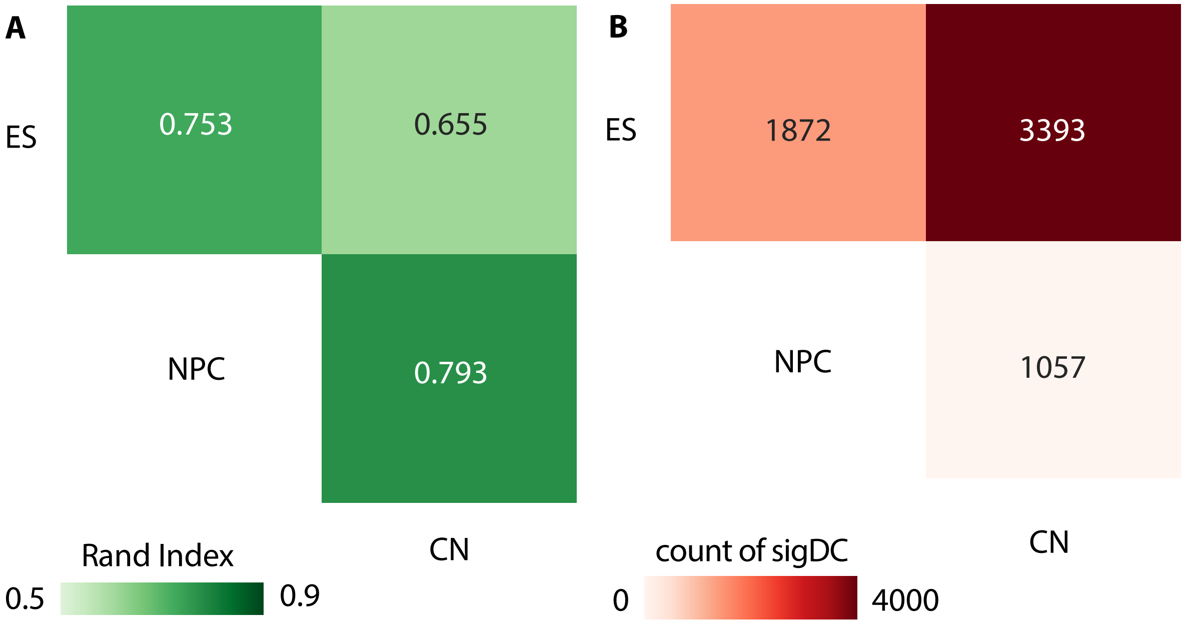

**Supp Figure 5.** Characterizing TGIF-DC clusters and sigDC on mouse neural differentiation data. **(A)** Similarity of cluster assignments for every pair of timepoints/states measured by Rand index. **(B)** Count of significantly differential compartmental regions (sigDC) for every pair of timepoints/states.

#### 1.6 Supp Figure 6

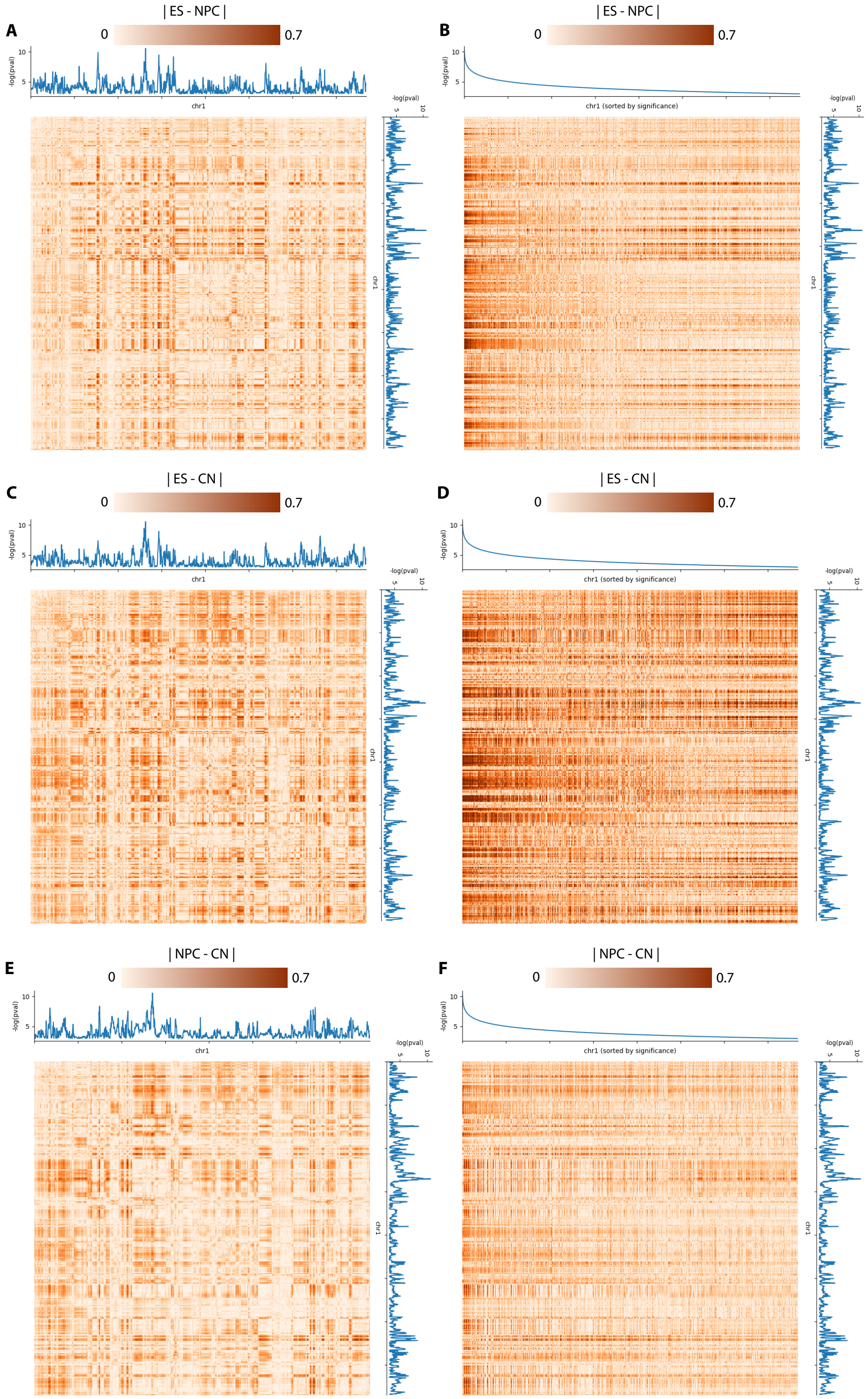

**(A)** Visualization of the difference in the input matrices (heatmap) and the significance of differences estimated with TGIF-DC (lineplot). Each row and column of the heatmap is a 100kb genomic region of chr1 and each entry in the heatmap = $\text{corr}(O/E)_{ES}$ - $\text{corr}(O/E)_{NPC}$. The lineplot shows -log(adjusted p-value) from TGIF-DC used for detecting significantly differential compartment regions between ES and NPC. **(B)** Same information as in **(A)**, but only the columns are sorted in descending significance. The sorting of regions by p-value highlights greater differences in count for regions with higher negative log p-values (high significance). **(C)** Same visualization as **(A)**, but for finding sigDC between ES and CN. **(D)** Same visualization as **(B)**, but for finding sigDC between ES and CN. **(E)** Same visualization as **(A)**, but for finding sigDC between NPC and CN. **(F)** Same visualization as **(B)**, but for finding sigDC between NPC and CN.

#### 1.7 Supp Figure 7

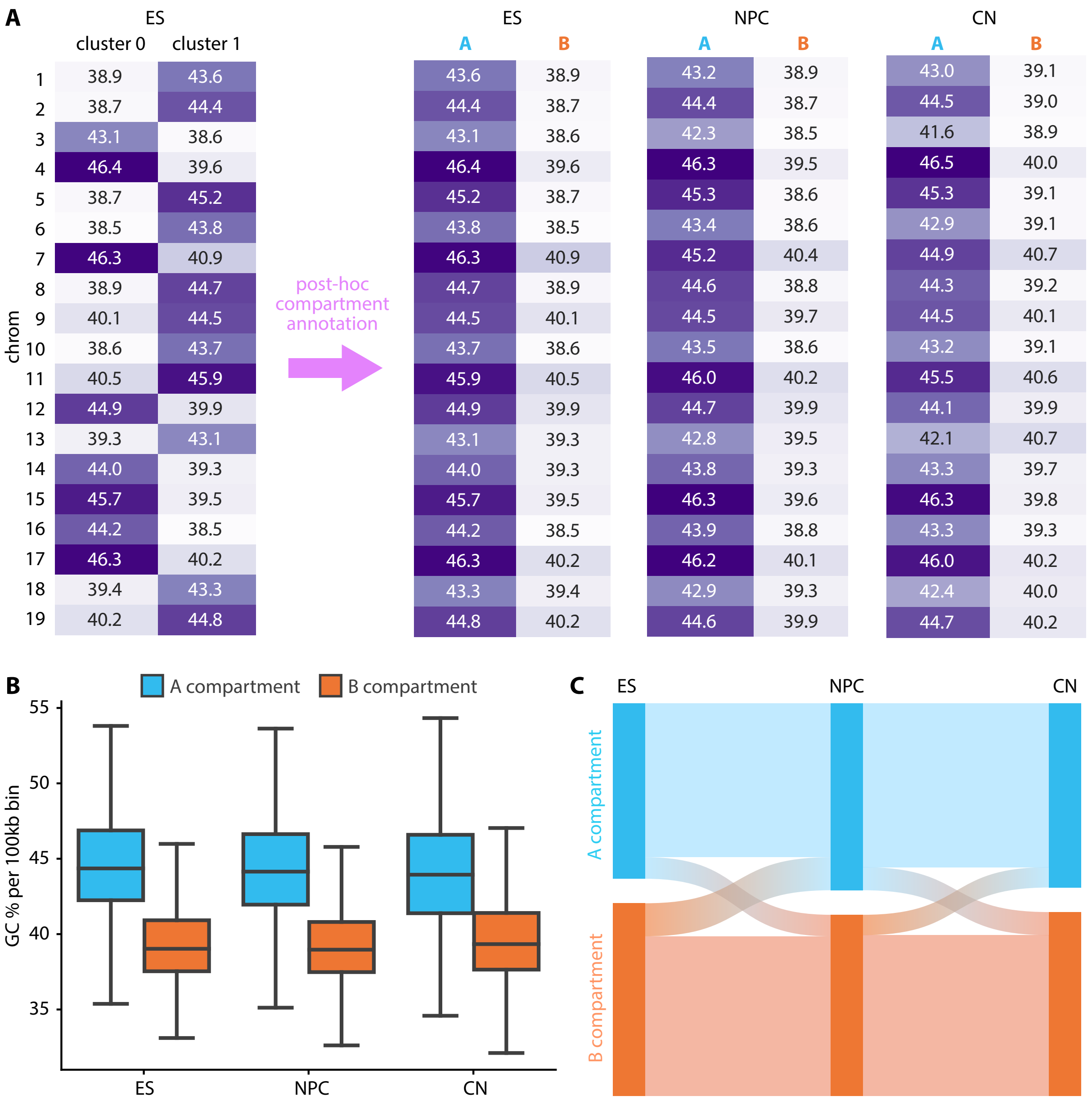

**Supp Figure 7.** Post-hoc annotation of TGIF-DC clusters into A and B compartments in mouse neural differentiation data. **(A)** Within each chromosome, the mean GC percentage is measured for regions in each TGIF-DC cluster for ES. Cluster with higher mean GC content is assigned to compartment A; the other to compartment B across all timepoints. **(B)** Genome-wide distribution of GC percentage in each 100kb bin by compartment assignment and timepoint. **(C)** Dynamic compartment assignment patterns for all genomic regions from ES to NPC to CN state.

#### 1.8 Supp Figure 8

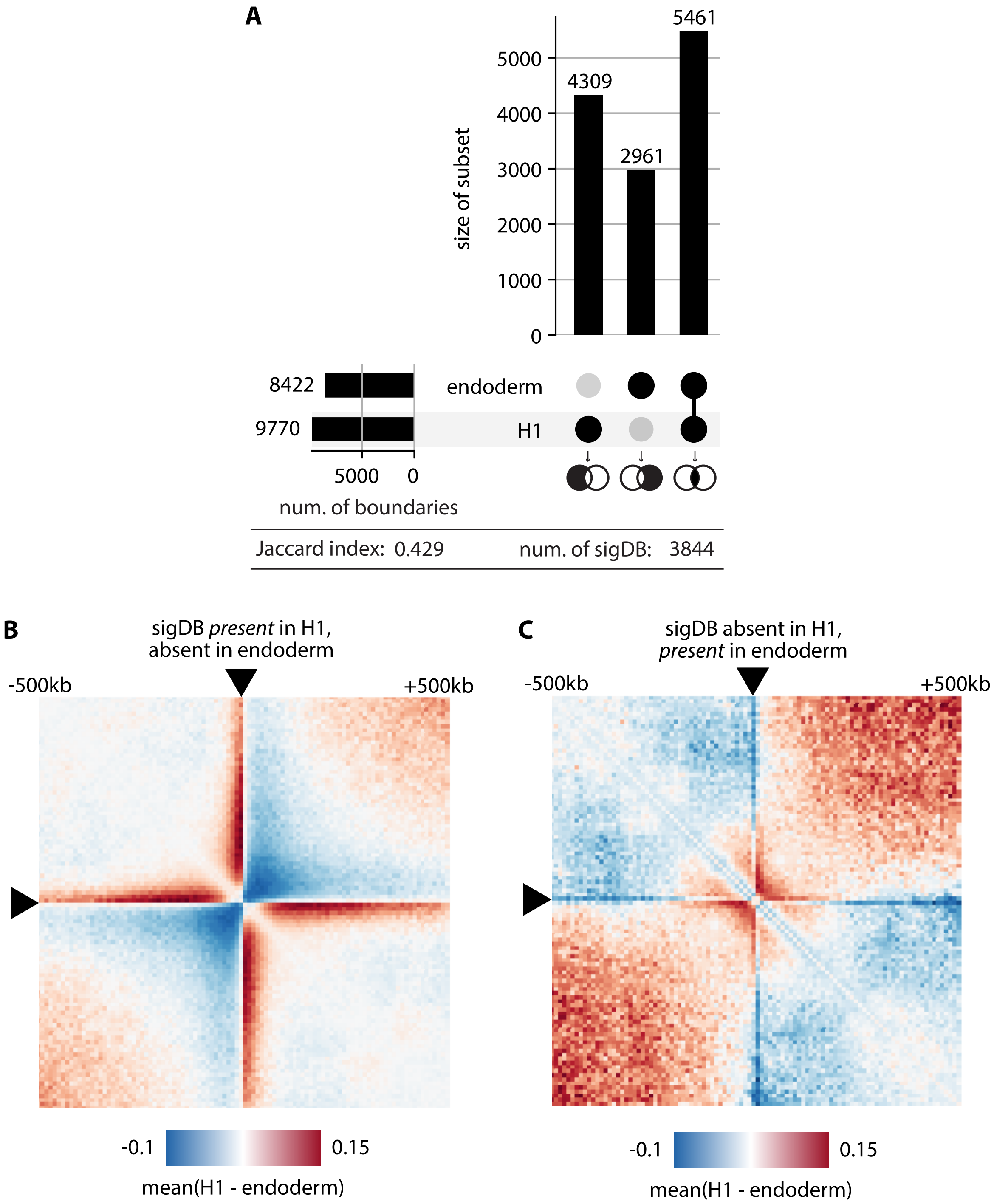

**Supp Figure 8.** Characterizing boundaries and sigDB in H1-endoderm differentiation. **(A)** Number of significant boundaries identified by TGIF-DB in H1 and differentiated endoderm. The vertical bars represent specific subsets only belonging to each category, i.e. boundaries unique to H1; those unique to endoderm; intersection of H1 and endoderm boundaries. The horizontal bars are total counts of boundaries identified in each state. Similarity of the boundary sets between H1 and endoderm is measured by Jaccard index. **(B)** Mean interaction count difference between H1 and endoderm near sigDB regions (surrounding 1MB window), present in H1 and absent in endoderm. To offset the depth difference between H1 and endoderm, interaction counts were first normalized to O/E matrices. **(C)** Mean interaction count difference between H1 and endoderm near sigDBs absent in H1 and present in endoderm.

#### 1.9 Supp Figure 9

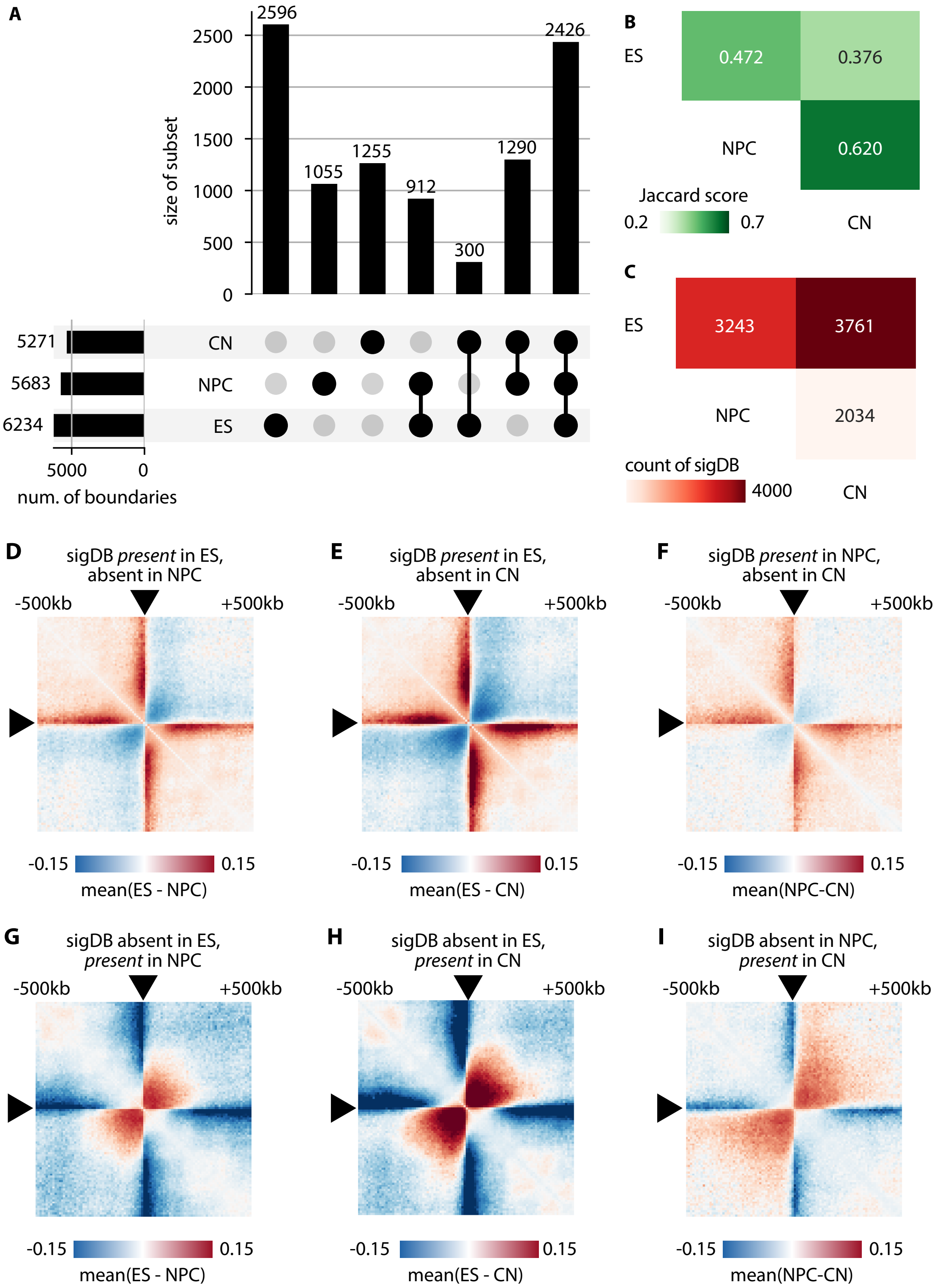

**Supp Figure 9.** Characterizing boundaries and sigDB in mouse neural differentiation. **(A)** Number of significant boundaries. The vertical bars represent specific subsets only belonging to each category, e.g. boundaries unique to ES, NPC, CN; intersection of ES and NPC boundaries. The horizontal bars are total counts of boundaries identified in each state. **(B)** Similarity of boundary sets between pairs of timepoints/states, measured by Jaccard index. **(C)** Count of significantly *differential* boundaries between pairs of timepoints/states. **(D)** Mean interaction count difference between ES and NPC near sigDB regions (the surrounding 1MB window), present in ES and absent in NPC. To offset the depth difference between ES and NPC, interaction counts were first normalized to O/E matrices. **(E)** Same visualization as in **(D)**, but for sigDBs present in ES but absent in CN. **(F)** Same visualization as in **(D)**, but for sigDBs resent in NPC but absent in CN. **(G)** Same visualization as in **(D)**, but for sigDBs absent in ES but present in NPC. **(H)** Same visualization as in **(D)**, but for sigDBs absent in ES but present in CN. **(I)** Same visualization as in **(D)**, but for sigDBs absent in NPC but present in CN.

#### 1.10 Supp Figure 10

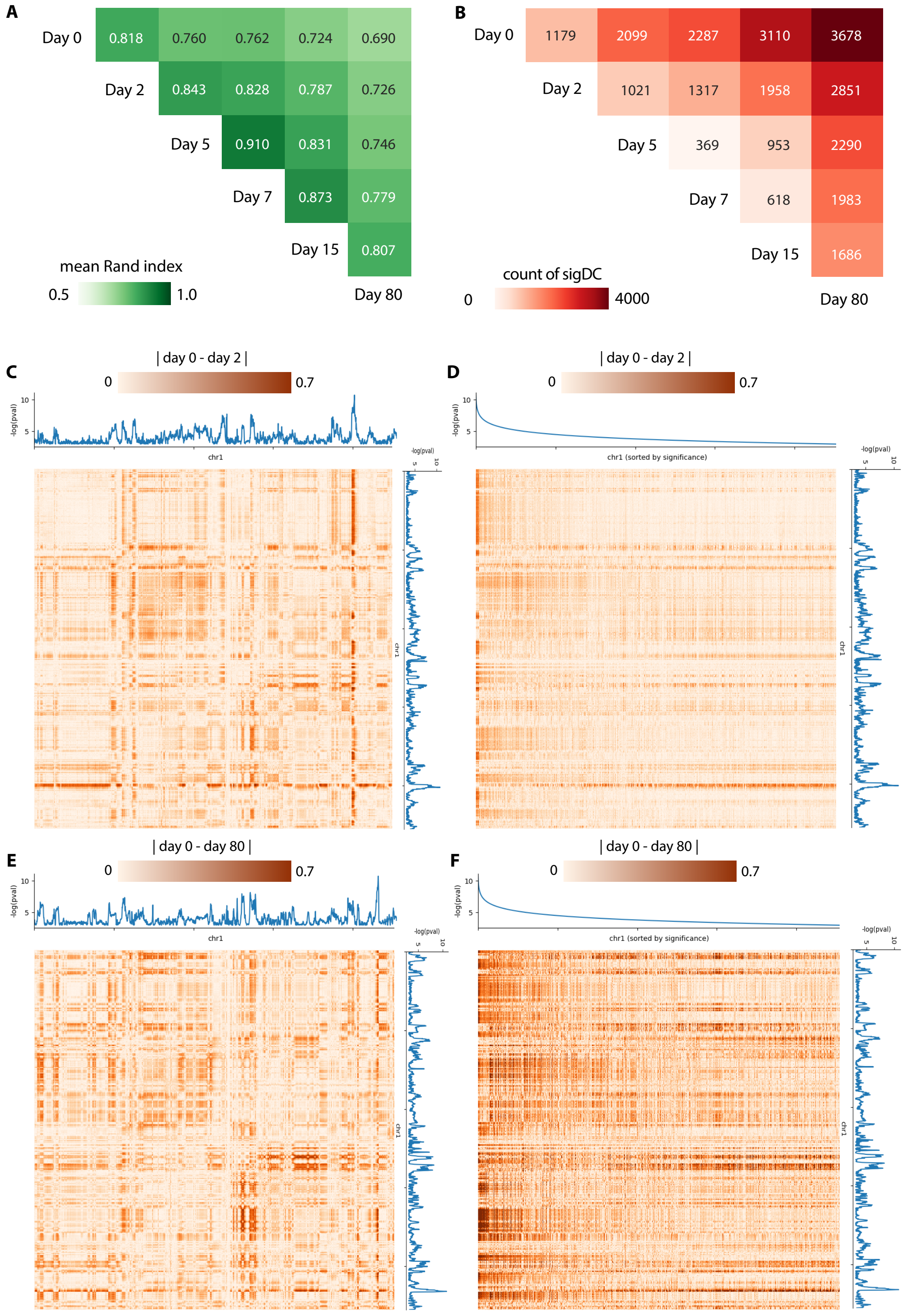

**Supp Figure 10.** Characterizing TGIF clusters and sigDC from applying TGIF-DC on cardiomyocyte differentiation data. **(A)** Similarity of compartment assignments for every pair of timepoints/states measured by Rand index. **(B)** Count of significantly differential compartmental regions (sigDC) for every pair of timepoints/states. **(C)** Visualization of the difference in the input matrices (heatmap) and the significance of differences estimated with TGIF-DC (lineplot). Each row and column of the heatmap is a 100kb genomic region of chr1 and each entry in the heatmap = $\text{corr(O/E)}_{\text{day 0}}$ - $\text{corr(O/E)}_{\text{day 2}}$. The lineplot shows -log(adjusted p-value) from TGIF-DC used for detecting significantly differential compartment regions between day 0 and day 2. **(D)** Same information as in **(C)**, but only the columns are sorted in descending significance. The sorting of regions by p-value highlights greater differences in count for regions with higher negative log p-values (high significance). **(E)** Same visualization as **(C)**, but for finding sigDC between day 0 and day 80. **(F)** Same visualization as **(D)**, but for finding sigDC between day 0 and day 80.

#### 1.11 Supp Figure 11

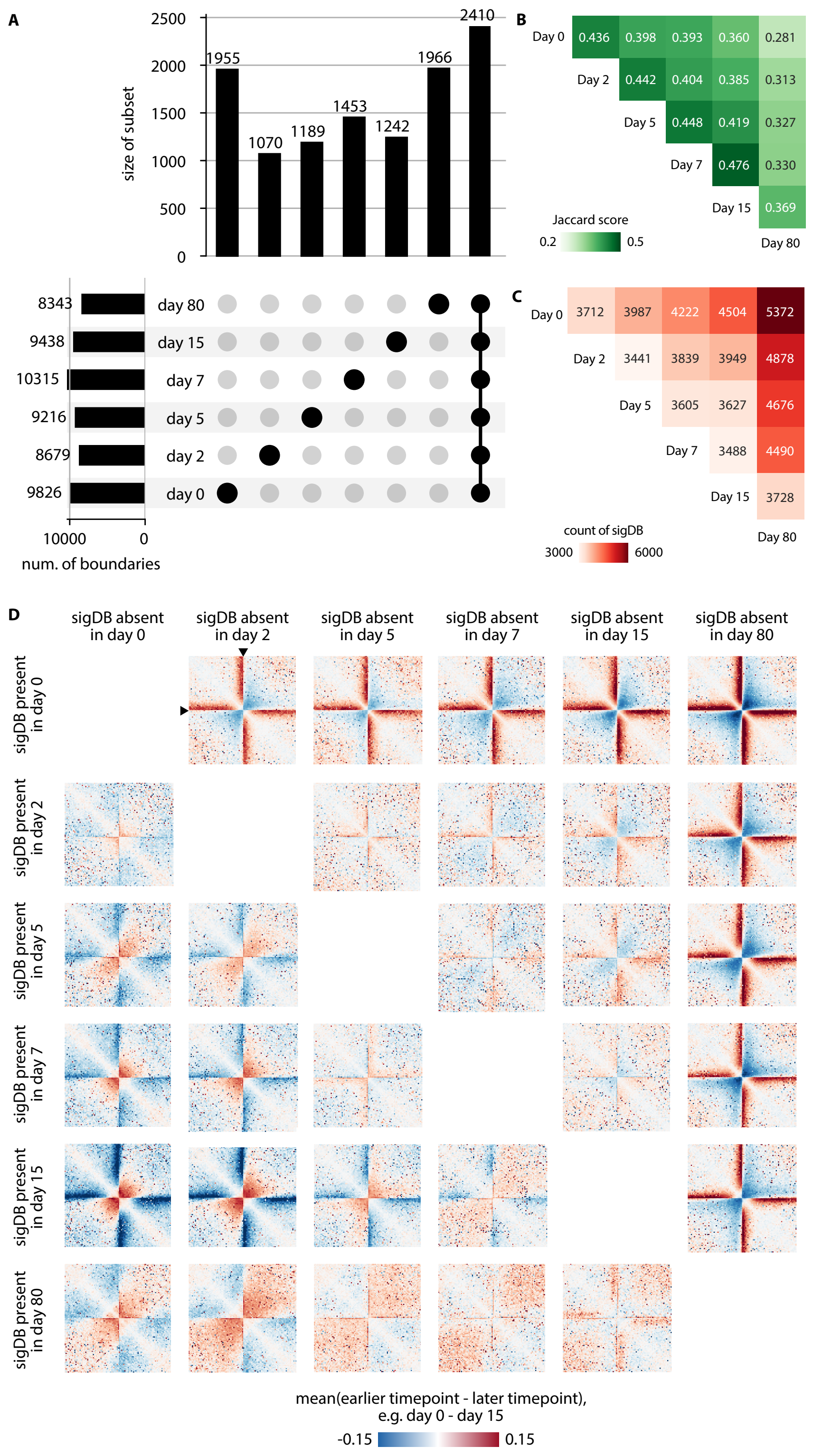

**Supp Figure 11.** Characterizing boundaries and sigDB in cardiomyocyte differentiation. **(A)** Number of significant boundaries. The vertical bars represent specific subsets only belonging to each category, e.g. boundaries unique to day 0, day 2, etc., intersection of day 0 and 2 boundaries. etc. The horizontal bars are total counts of boundaries identified in each timepoint. **(B)** Similarity of boundary sets between pairs of timepoints/states, measured by Jaccard index. **(C)** Count of significantly *differential* boundaries between pairs of timepoints/states. **(D)** Mean interaction count difference between two timepoints near sigDB regions (the surrounding 1MB window). Each row of the grid represents the timepoint in which the boundary is present and each column the timepoint in which it is absent. To offset the depth difference between the two timepoints, interaction counts were first normalized to O/E matrices. Each heatmap is the count different matrix where the counts from the earlier timepoint is subtracted by those from the later timepoint in the pair as indicated.

#### 1.12 Supp Figure 12

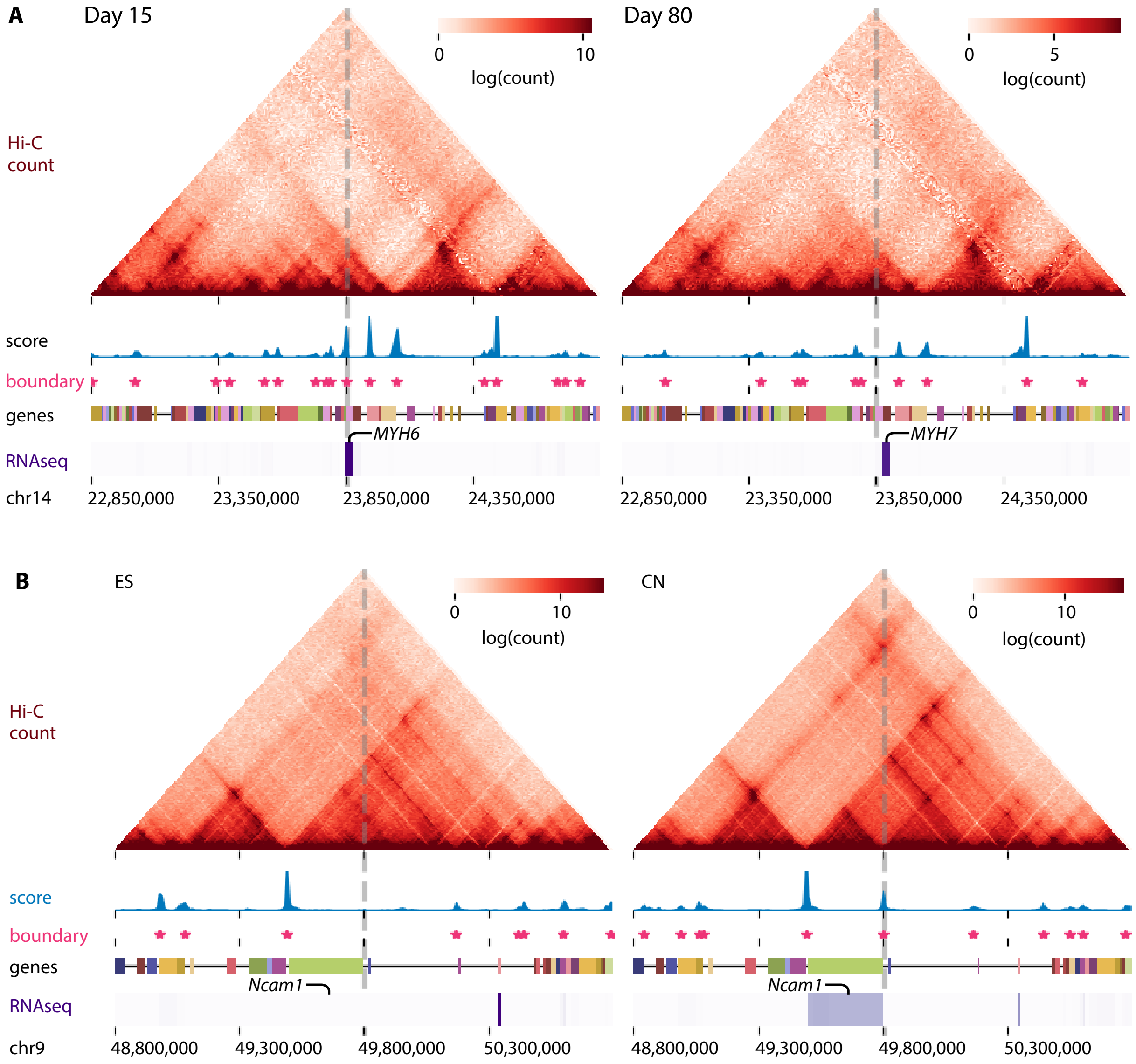

**Supp Figure 12.** Examples of differentially expression (DE) gene near significantly differential boundary (sigDB). **(A)** A highly ranked sigDB (loci marked with dotted vertical line) close to a DE gene, *MYH6*. Shown are the Hi-C interaction matrices for the ES and CN stages of mouse neural differentiation, the boundary score (blue), significant boundary (asterisk), a gene track and the expression heatmap from RNA-seq data. Differential expression of *MYH6* near a sigDB present in day 15 of cardiomyocyte differentiation but absent in day 80 is shown. **(B)** A sigDB close to a DE gene, *Ncam1*. Differential expression of *Ncam1* near the sigDB that is absent in ES state but appears in CN is shown.

#### 1.13 Supp Figure 13

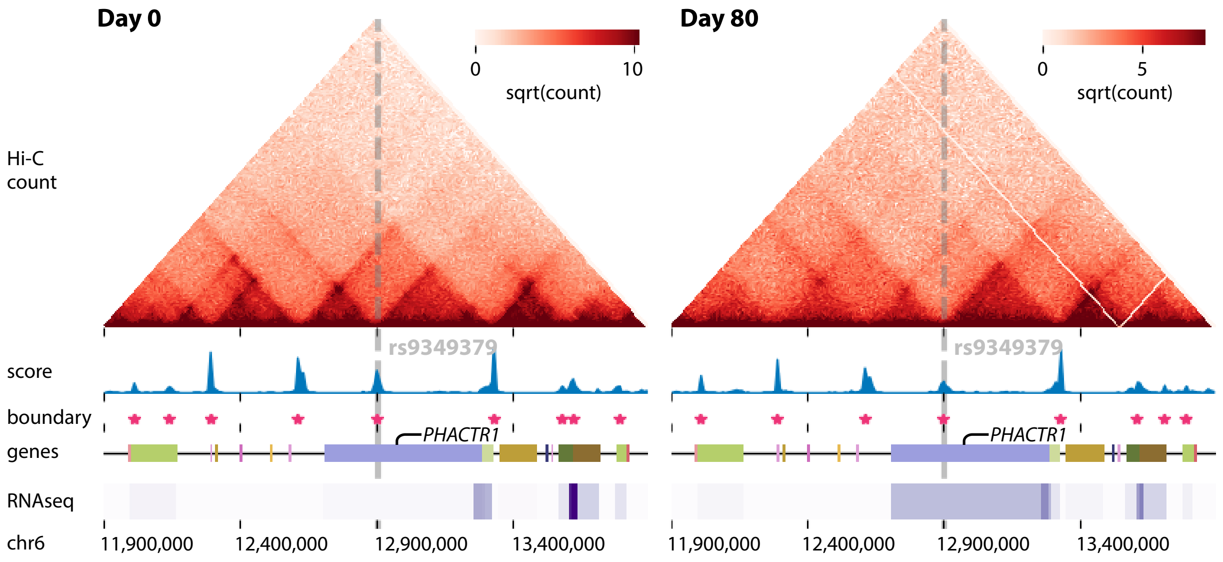

**Supp Figure 13.** Persistent TGIF boundaries identified from human cardiomyocyte differentiation containing the cardiovascular disease associated GWAS SNP rs9349379.

#### 1.14 Supp Figure 14

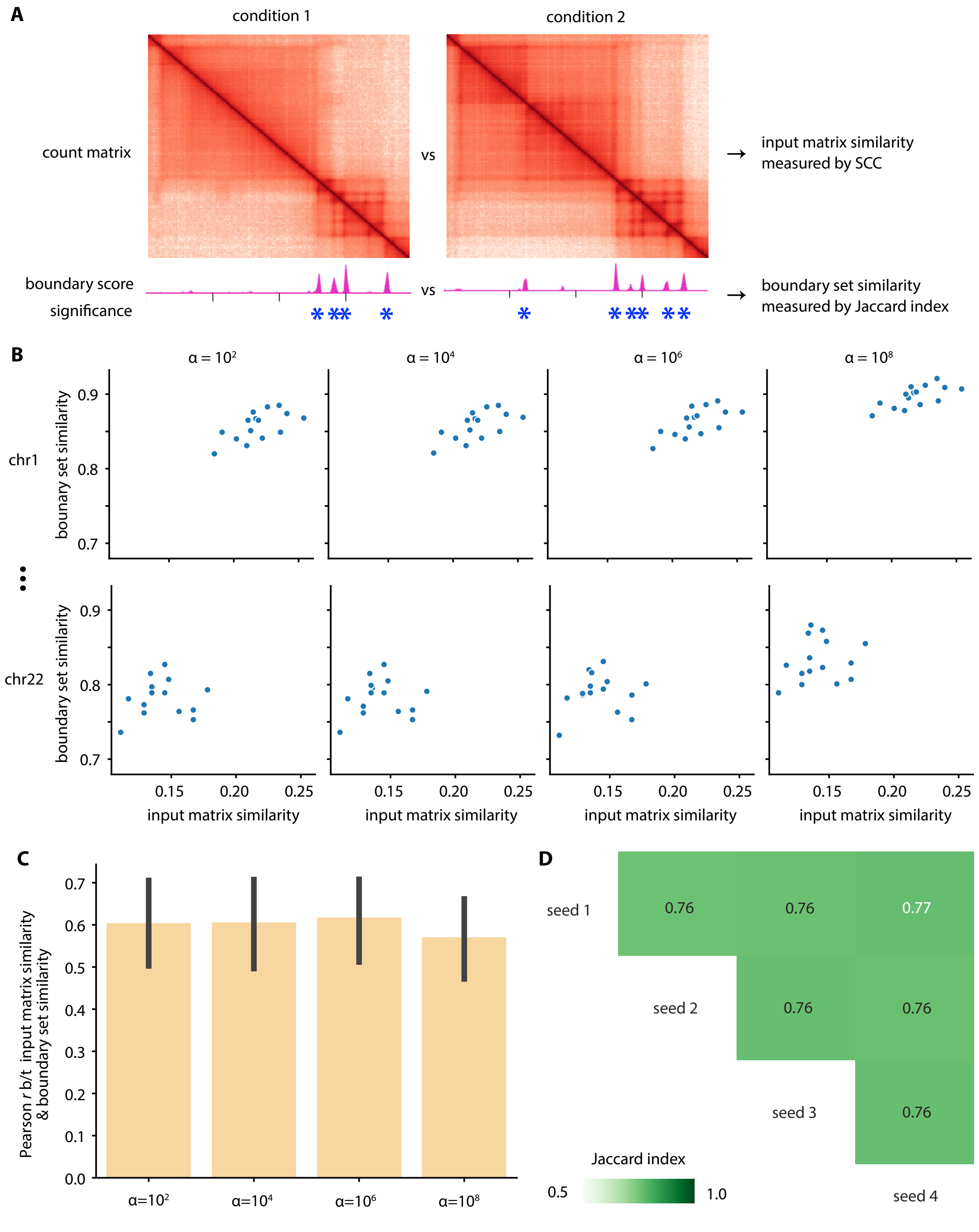

**Supp Figure 14.** Hyperparameter $\alpha$ selection for TGIF-DB. **(A)** The similarity between the input matrices is measured by SCC and the output boundary set agreement measured by Jaccard index. **(B)** Plotting input matrix similarity vs output boundary set agreement for each $\alpha$ and each chromosome. Each dot represents one pairwise comparison. **(C)** Correlation between the input matrix similarity and the output boundary set agreement for different values of $\alpha$. **(D)** Similarity of boundary sets from different random initialization seeds (with $\alpha = 10^6$), measured by Jaccard index.

#### 1.15 Supp Figure 15

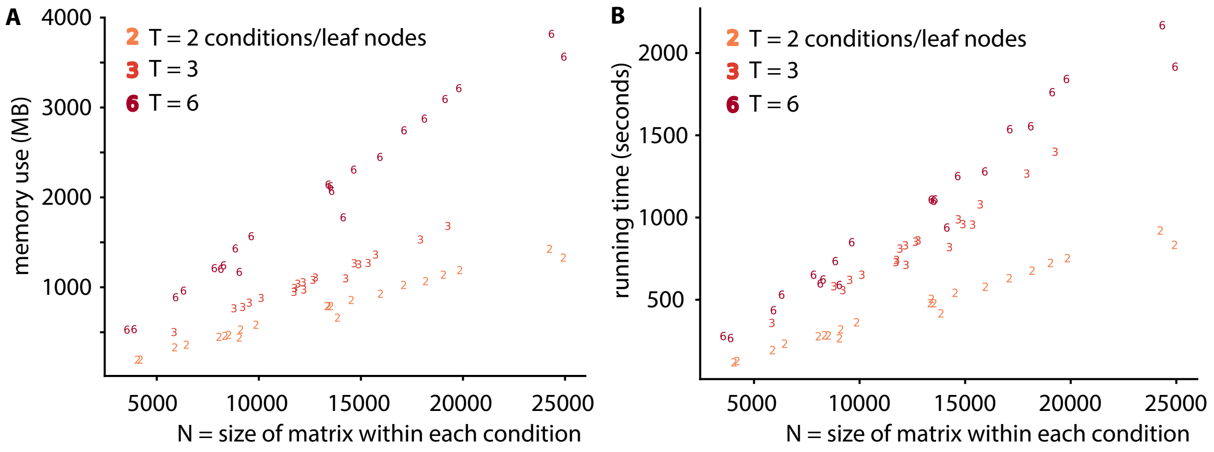

**Supp Figure 15.** Resource use by TGIF-DB. **(A)** Size of each input matrix (100kb resolution) vs memory use in MB. **(B)** Size of each input matrix vs running time in seconds. $T$ = the number of conditions/states/timepoints in a given run of TGIF-DB.

#### 1.16 Supp Figure 16

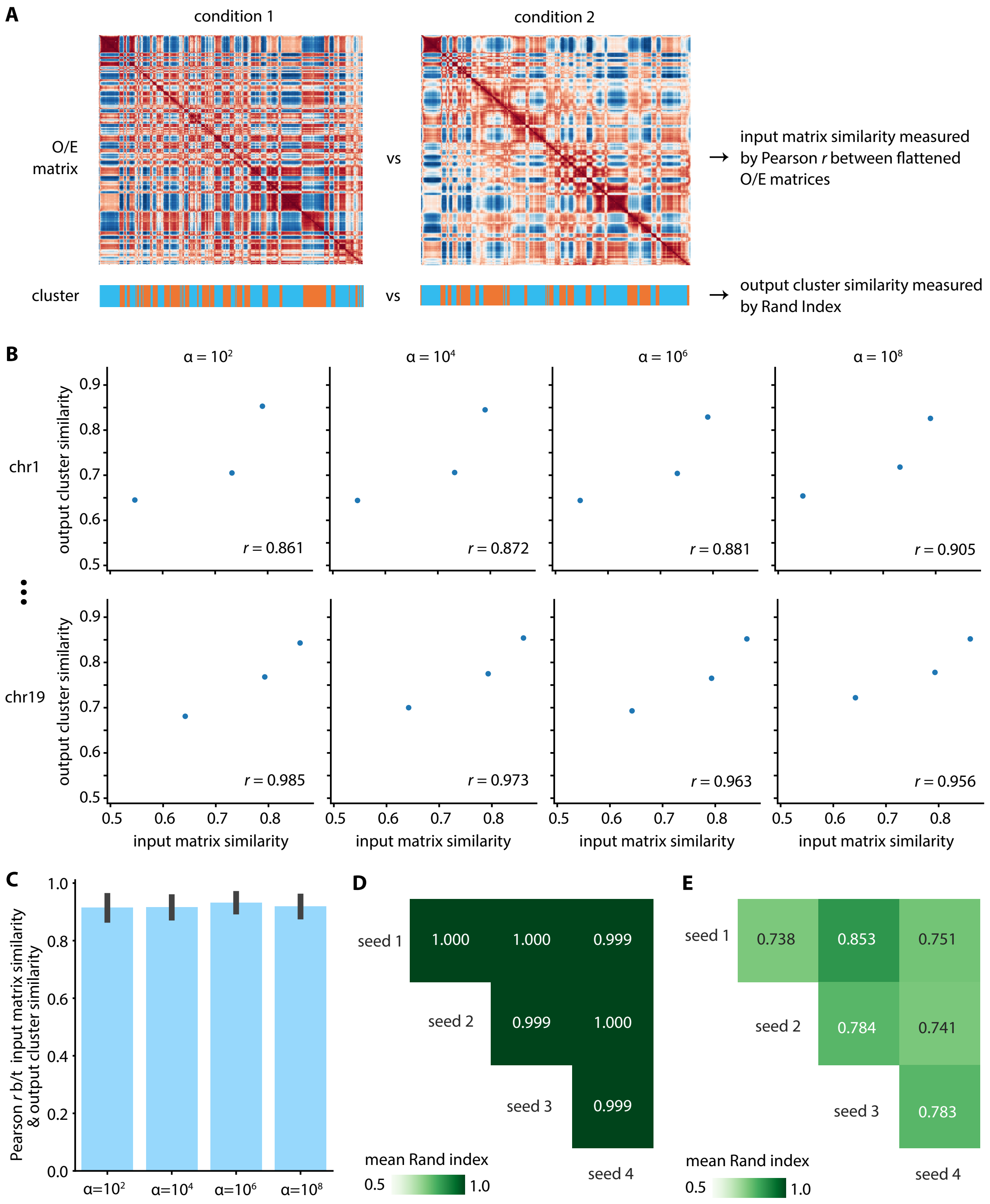

**Supp Figure 16.** Hyperparameter $\alpha$ selection for TGIF-DC. **(A)** The similarity between the O/E count matrices is measured by correlation of the flattened matrices the output cluster similarity measured by Rand index. **(B)** Plotting input matrix similarity vs output cluster similarity for each $\alpha$ and each chromosome. Each dot represents one pairwise comparison. **(C)** Correlation between the input matrix similarity and the output cluster similarity for different values of $\alpha$. **(D)** Similarity of cluster assignments from different random initialization seeds (with $\alpha = 10^4$), measured by Rand index.

#### 1.17 Supp Figure 17

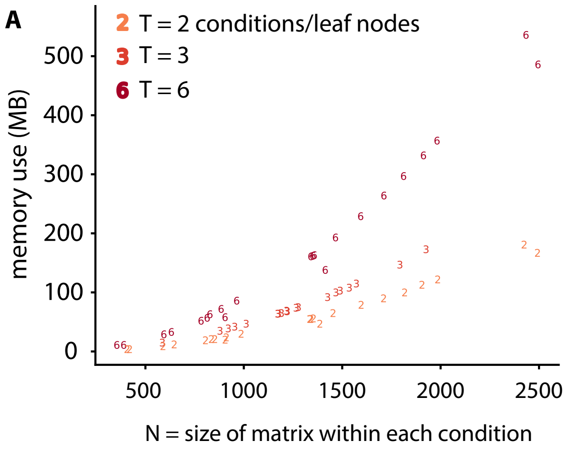

**Supp Figure 17.** Resource use by TGIF-DC. **(A)** Size of each input matrix (100kb resolution) vs memory use in MB. **(B)** Size of each input matrix vs running time in seconds. $T$ = the number of conditions/states/timepoints in a given run of TGIF-DC.

#### 1.18 Supp Figure 18

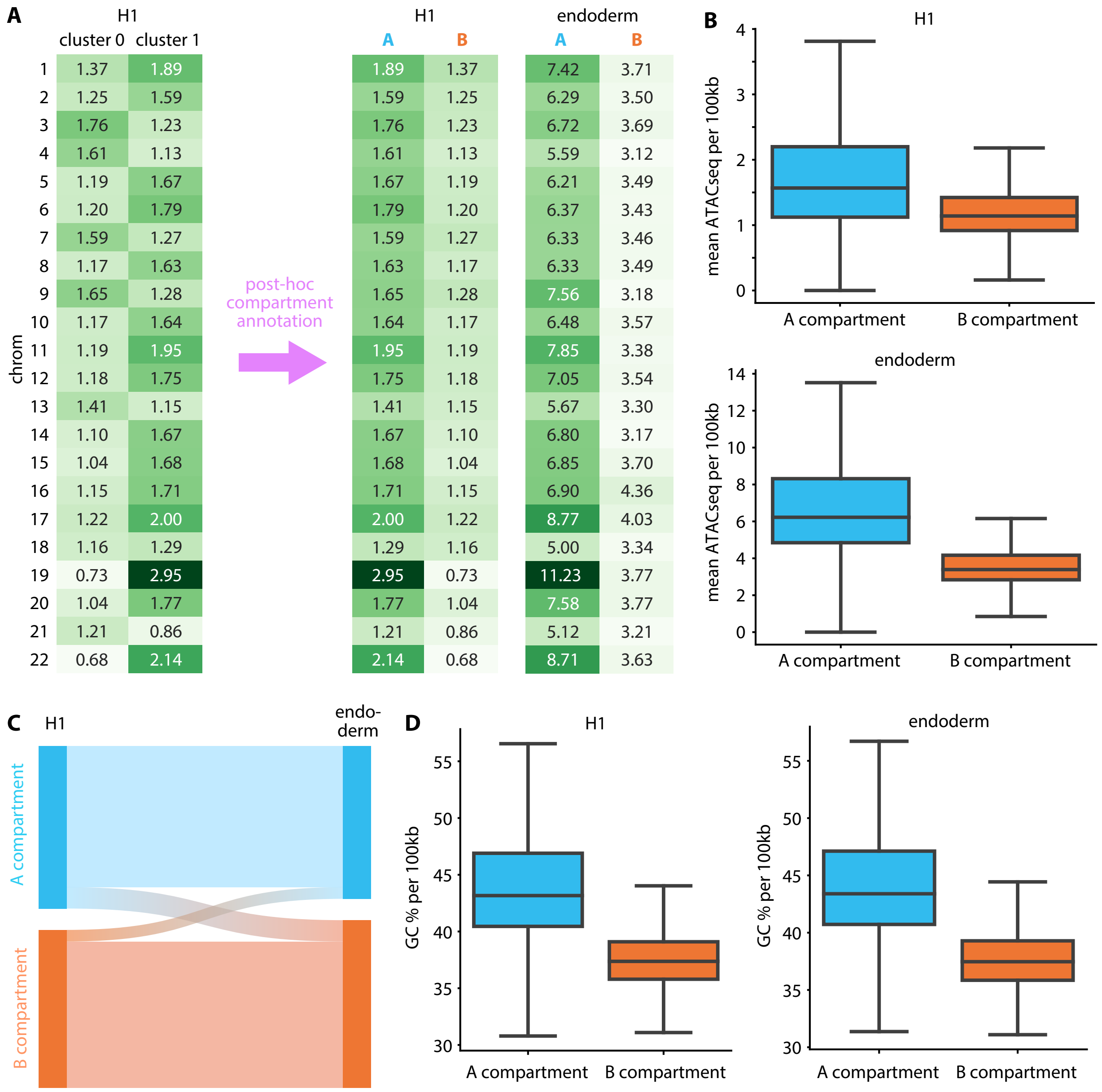

**Supp Figure 18.** Post-hoc annotation of TGIF-DC clusters into A and B compartments in H1-endoderm dataset using accessibility. **(A)** For each chromosome, mean ATACseq signal in each 100kb bin is measured for each TGIF-DC cluster in H1. The cluster with higher mean ATACseq signal is annotated as A compartment; the other cluster B compartment. **(B)** Genome-wide mean ATACseq signal distribution by compartment in H1 (top) and endoderm (bottom). **(C)** Dynamic compartment assignment pattern from H1 to endoderm. **(D)** Genome-wide mean GC content per 100kb bin by compartment in H1 and endoderm.

#### 1.19 Supp Figure 19

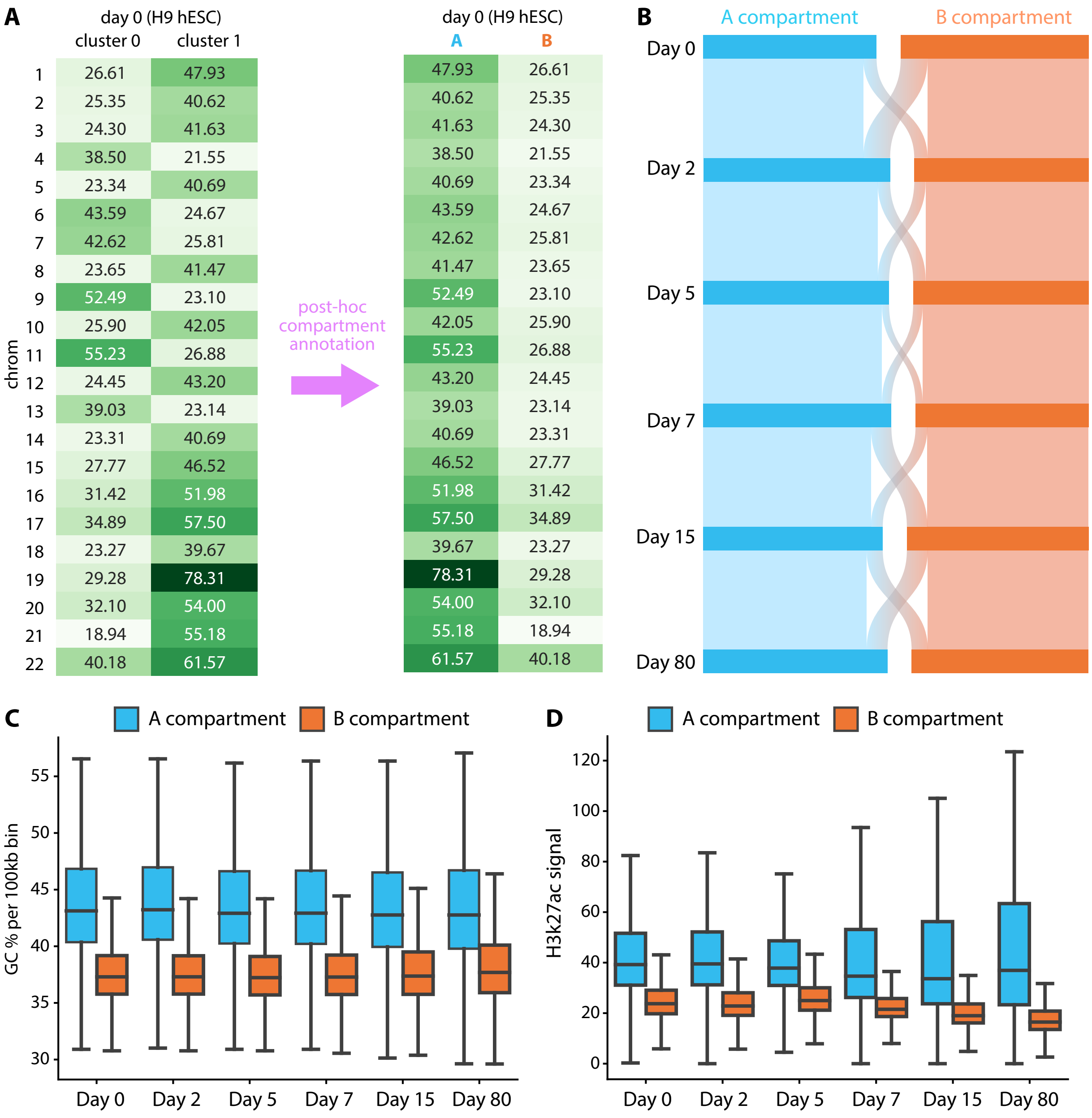

| chrom | cluster 0 | cluster 1 |
|---|---|---|
| 1 | 26.61 | 47.93 |
| 2 | 25.35 | 40.62 |
| 3 | 24.30 | 41.63 |
| 4 | 38.50 | 21.55 |
| 5 | 23.34 | 40.69 |
| 6 | 43.59 | 24.67 |
| 7 | 42.62 | 25.81 |
| 8 | 23.65 | 41.47 |
| 9 | 52.49 | 23.10 |
| 10 | 25.90 | 42.05 |
| 11 | 55.23 | 26.88 |
| 12 | 24.45 | 43.20 |
| 13 | 39.03 | 23.14 |
| 14 | 23.31 | 40.69 |
| 15 | 27.77 | 46.52 |
| 16 | 31.42 | 51.98 |
| 17 | 34.89 | 57.50 |
| 18 | 23.27 | 39.67 |
| 19 | 29.28 | 78.31 |
| 20 | 32.10 | 54.00 |
| 21 | 18.94 | 55.18 |
| 22 | 40.18 | 61.57 |

| A | B |
|---|---|
| 47.93 | 26.61 |
| 40.62 | 25.35 |
| 41.63 | 24.30 |
| 38.50 | 21.55 |
| 40.69 | 23.34 |
| 43.59 | 24.67 |
| 42.62 | 25.81 |
| 41.47 | 23.65 |
| 52.49 | 23.10 |
| 42.05 | 25.90 |
| 55.23 | 26.88 |
| 43.20 | 24.45 |
| 39.03 | 23.14 |
| 40.69 | 23.31 |
| 46.52 | 27.77 |
| 51.98 | 31.42 |
| 57.50 | 34.89 |
| 39.67 | 23.27 |
| 78.31 | 29.28 |
| 54.00 | 32.10 |
| 55.18 | 18.94 |
| 61.57 | 40.18 |

**Supp Figure 19.** Post-hoc annotation of TGIF-DC clusters into A and B compartments in cardiomyocyte differentiation dataset using accessibility. **(A)** For each chromosome, mean DNaseq signal in each 100kb bin is measured for each TGIF-DC cluster in day 0. The cluster with higher mean DNaseq signal is annotated as A compartment; the other cluster B compartment. **(B)** Dynamic compartment assignment pattern from day 0 to day 80. **(C)** Genome-wide mean GC content per 100kb bin by compartment across all timepoints. **(D)** Genome-wide mean H3k27ac signal distribution by compartment across all timepoints.

### 2 Supplementary Table

#### 2.1 Supp Table 1

4D Nucleome accession numbers for H1 and endoderm data.

#### 2.2 Supp Table 2

Data sources and GEO accession numbers for mouse neural differentiation data.

#### 2.3 Supp Table 3

GEO and ENCODE accession numbers for cardiomyocyte differentiation data.

#### 2.4 Supp Table 4

4D Nucleome accession number for GM12878.

#### 2.5 Supp Table 5

Differential expression fold enrichment analysis on H1-endoderm dataset. Each row represents one fold enrichment hypergeometric test.

#### 2.6 Supp Table 6

Differential expression fold enrichment analysis on mouse neural differentiation dataset. Each row represents one fold enrichment hypergeometric test.

#### 2.7 Supp Table 7

Differential expression fold enrichment analysis on cardiomyocyte dataset. Each row represents one fold enrichment hypergeometric test.

#### 2.8 Supp Table 8

GO enrichment analysis on cadiomyocyte differentiation dataset. Each entry is the negative log p-value of GO biological processes for two subsets of differentially expressed (DE) genes: DE genes close to sigDB, and DE genes not close to sigDB. Each column is a pair of timepoints used to determine DB and DE genes. Each row is a GO term (only GO terms with p-value $<$ 1e-5 in at least one pair of timepoints is listed). Non-blank entries are the negative log p-value of the GO term enrichment.

#### 2.9 Supp Table 9

GO enrichment analysis on H1-endoderm dataset. Each entry is the negative log p-value of GO biological processes for two subsets of differentially expressed (DE) genes: DE genes close to sigDB, and DE genes not close to sigDB. Each column is a pair of timepoints used to determine DB and DE genes. Each row is a GO term (only GO terms with p-value $<$ 1e-5 in at least one pair of timepoints is listed). Non-blank entries are the negative log p-value of the GO term enrichment.

#### 2.10 Supp Table 10

GO enrichment analysis on mouse neural differentiation dataset. Each entry is the negative log p-value of GO biological processes for two subsets of differentially expressed (DE) genes: DE genes close to sigDB, and DE genes not close to sigDB. Each column is a pair of timepoints used to determine DB and DE genes. Each row is a GO term (only GO terms with p-value $< 1e\text{-}5$ in at least one pair of timepoints is listed). Non-blank entries are the negative log p-value of the GO term enrichment.

#### 3 Supplementary Methods: Block Coordinate Descent for TGIF

Below we walk through the derivation for the Block Coordinate Descent (BCD) optimization and update rules for Tree-Guided Integrated Factorization (TGIF).

##### 3.1 Notation and objective

Given $t \in \{1, \dots, T\}$ tasks, each with input matrix $X^{(t)} \in \mathbb{R}^{n_t \times m}$, related to each other in a task hierarchy/tree with a set of nodes $c \in \{r\} \cup \mathcal{B} \cup \mathcal{T}$ where $r$ is the root node, $\mathcal{B}$ a set of internal (or branch) nodes $b \in \mathcal{B}$, and $\mathcal{T}$ a set of the task-specific leaf nodes, the objective is:

$$O = \sum_{t=1}^{T} \left\| X^{(t)} - U^{(t)} V^{(t)\top} \right\|_{\mathrm{F}}^2 + \alpha \sum_{c} \left\| V^{(c)} - V^{Pa(c)} \right\|_{\mathrm{F}}^2 \tag{1}$$

where $U^{(t)} \in \mathbb{R}^{n_t \times k}$, $V^{(\cdot)} \in \mathbb{R}^{m \times k}$, $k \ll n, m$. The regularization term will:

1. constrain a task-specific latent feature factor $V^{(t)}$ in a leaf node of the task hierarchy to be similar to $V^{\mathrm{Pa}(t)}$ in its parent node;
2. constrain an internal node's latent feature factor $V^{(b)}$ to be similar to its direct child nodes' $V^{(c)}$ and and its parent node's $V^{Pa(b)}$; and
3. constrain the root node's latent feature factor $V^{(r)}$ to be similar to all of its direct child nodes' $V^{(c)}$s.

##### 3.2 Breaking down to task-level and column-level subproblems

The objective can be re-written as:

$$O = \sum_{t=1}^{T} \left\| X^{(t)} - \sum_{k} u_k^{(t)} v_k^{(t)\top} \right\|_{\mathrm{F}}^2 + \alpha \sum_{c} \sum_{k} \left\| v_k^{(c)} - v_k^{Pa(c)} \right\|_2^2 \tag{2}$$

Where $u_k^{(t)} \in \mathbb{R}^{n_t}$ is the $k$th column vector of $U^{(t)}$ and $v_k^{(t)} \in \mathbb{R}^m$ is the $k$th column vector of $V^{(t)}$. Now we 'pull out' terms involving the $k$th column in all factors:

$$O = \sum_{t=1}^{T} \left\| X^{(t)} - u_k^{(t)} v_k^{(t)\top} - \sum_{j \neq k} u_j^{(t)} v_j^{(t)\top} \right\|_{\mathrm{F}}^2 + \alpha \sum_{c} \left( \left\| v_k^{(c)} - u_k^{Pa(c)} \right\|_2^2 + \sum_{j \neq k} \left\| v_j^{(c)} - v_j^{Pa(c)} \right\|_2^2 \right) \tag{3}$$

Now we will substitute with $R_k^{(t)} = X^{(t)} - \sum_{j \neq k} u_j^{(t)} v_j^{(t)\top}$:

$$O = \sum_{t=1}^{T} \left\| R_k^{(t)} - u_k^{(t)} v_k^{(t)\top} \right\|_{\mathrm{F}}^2 + \alpha \sum_{c} \left\| v_k^{(c)} - v_k^{Pa(c)} \right\|_2^2 + \alpha \sum_{c} \sum_{j \neq k} \left\| v_j^{(c)} - v_j^{Pa(c)} \right\|_2^2 \tag{4}$$

We can now attempt to optimize $u_k^{(t)}$ and $v_k^{(\cdot)}$, fixing all other parameters to be constant.

##### 3.2.1 Optimize $v_k^{(t)}$

To find $v_k^{(t)}$ for each leaf node task $t$ that minimizes the objective, we find the derivative of the objective with respect to $v_k^{(t)}$ and set it to 0, then solve. First we expand the objective into matrix multiplications:

$$O = \left\| R_k^{(t)} - u_k^{(t)} v_k^{(t)\top} \right\|_{\text{F}}^2 + \alpha \left\| v_k^{(t)} - v_k^{Pa(t)} \right\|_2^2 + C \tag{5}$$

$$= \operatorname{Tr}\left[ \left( R_k^{(t)} - u_k^{(t)} v_k^{(t)\top} \right)^\top \left( R_k^{(t)} - u_k^{(t)} v_k^{(t)\top} \right) \right] + \alpha \left( v_k^{(t)} - v_k^{Pa(t)} \right)^\top \left( v_k^{(t)} - v_k^{Pa(t)} \right) + C \tag{6}$$

Here $C$ subsumes all elements of the objective that does not involve $v_k^{(t)}$ (including terms involving tasks other than $t$), since they will be zeroed out when the derivative is taken with respect to $v_k^{(t)}$. Now we keep expanding:

$$O = \operatorname{Tr}\left[ R_k^{(t)\top} R_k^{(t)} - 2 R_k^{(t)\top} u_k^{(t)} v_k^{(t)\top} + \left( u_k^{(t)} v_k^{(t)\top} \right)^\top \left( u_k^{(t)} v_k^{(t)\top} \right) \right] \tag{7}$$

$$+ \alpha \left( v_k^{(t)\top} v_k^{(t)} - 2 v_k^{(t)\top} v_k^{Pa(t)} + v_k^{Pa(t)\top} v_k^{Pa(t)} \right) + C \tag{8}$$

$$= \operatorname{Tr}\left( R_k^{(t)\top} R_k^{(t)} \right) - 2 \operatorname{Tr}\left( R_k^{(t)\top} u_k^{(t)} v_k^{(t)\top} \right) + \operatorname{Tr}\left( v_k^{(t)} u_k^{(t)\top} u_k^{(t)} v_k^{(t)\top} \right) \tag{9}$$

$$+ \alpha v_k^{(t)\top} v_k^{(t)} - 2\alpha v_k^{(t)\top} v_k^{Pa(t)} + \alpha v_k^{Pa(t)\top} v_k^{Pa(t)} + C \tag{10}$$

$$= \operatorname{Tr}\left( R_k^{(t)\top} R_k^{(t)} \right) - 2 \left( R_k^{(t)\top} u_k^{(t)} \right)^\top v_k^{(t)} + \left( u_k^{(t)\top} u_k^{(t)} \right) \left( v_k^{(t)\top} v_k^{(t)} \right) \tag{11}$$

$$+ \alpha v_k^{(t)\top} v_k^{(t)} - 2\alpha v_k^{(t)\top} v_k^{Pa(t)} + \alpha v_k^{Pa(t)\top} v_k^{Pa(t)} + C \tag{12}$$

Now we take the derivative of $O$ w.r.t. $v_k^{(t)}$:

$$\frac{\partial O}{\partial v_k^{(t)}} = 0 - 2 R_k^{(t)\top} u_k^{(t)} + 2 v_k^{(t)} u_k^{(t)\top} u_k^{(t)} + 2\alpha v_k^{(t)} - 2\alpha v_k^{Pa(t)} + 0 + 0 \tag{13}$$

$$0 = -R_k^{(t)\top} u_k^{(t)} + \left( u_k^{(t)\top} u_k^{(t)} + \alpha \right) v_k^{(t)} - \alpha v_k^{Pa(t)} \tag{14}$$

$$v_k^{(t)} = \frac{R_k^{(t)\top} u_k^{(t)} + \alpha v_k^{Pa(t)}}{\left\| u_k^{(t)} \right\|_2^2 + \alpha} \tag{15}$$

With the non-negativity constraint $v_k^{(t)} \geq 0$, we want $R_k^{(t)\top} u_k^{(t)} + \alpha v_k^{Pa(t)} \geq 0$, because if $R_k^{(t)\top} u_k^{(t)} + \alpha v_k^{Pa(t)} < 0$, $O$ will increase in (11) and (12). So the finalized update rule is:

$$v_k^{(t)} = \frac{\left[ R_k^{(t)\top} u_k^{(t)} + \alpha v_k^{Pa(t)} \right]_+}{\left\| u_k^{(t)} \right\|_2^2 + \alpha} \tag{16}$$

##### 3.2.2 Optimize $u_k^{(t)}$

We can derive the update rule for $u_k^{(t)}$ in leaf node task $t$ similarly but much more simply. From (12), we take the derivative of $O_t$ with respect to $u_k^{(t)}$; all regularization terms will zero out since they do not involve

$u_k^{(t)}$. Hence the final update rule for $u_k^{(t)}$ is:

$$u_k^{(t)} = \frac{\left[R_k^{(t)} v_k^{(t)}\right]_+}{\left\|v_k^{(t)}\right\|_2^2} \tag{17}$$

##### 3.2.3 Optimize $v_k^{(r)}$

For the overall consensus factor in the root of the task hierarchy, $v_k^{(r)}$, we can again ignore terms that do not involve $v_k^{(r)}$ in the objective (4). Note that we're going to collect the terms involving nodes $c$ whose parent is the root node, i.e. Pa$(c) = r$:

$$\begin{aligned}
O &= \alpha \sum_{c \in \text{Child}(r)} \left\|v_k^{(c)} - v_k^{(r)}\right\|_2^2 + C && (18)\\
&= \alpha \sum_{c \in \text{Child}(r)} \left(v_k^{(c)} - v_k^{(r)}\right)^\top \left(v_k^{(c)} - v_k^{(r)}\right) + C && (19)\\
&= \alpha \sum_{c \in \text{Child}(r)} \left[v_k^{(c)\top} v_k^{(c)} - 2 v_k^{(c)\top} v_k^{(r)} + v_k^{(r)\top} v_k^{(r)}\right] + C && (20)\\
&= C - \sum_{c \in \text{Child}(r)} 2\alpha v_k^{(c)\top} v_k^{(r)} + \sum_{c \in \text{Child}(r)} \alpha v_k^{(r)\top} v_k^{(r)} && (21)
\end{aligned}$$

Now we take the derivative, set to 0, and solve:

$$\begin{aligned}
\frac{\partial O}{\partial v_k^{(r)}} &= 0 - \sum_{c \in \text{Child}(r)} 2\alpha v_k^{(c)} + \sum_{c \in \text{Child}(r)} 2\alpha v_k^{(r)} && (22)\\
0 &= - \sum_{c \in \text{Child}(r)} v_k^{(c)} + |\text{Child}(r)| \cdot v_k^{(r)} && (23)\\
v_k^{(r)} &= \frac{\sum_{c \in \text{Child}(r)} v_k^{(c)}}{|\text{Child}(r)|} && (24)
\end{aligned}$$

where $|\text{Child}(r)|$ is the number of direct child nodes of the root node $r$.

##### 3.2.4 Optimize $v_k^{(b)}$

For the latent feature factor in an internal/branch node of the task hierarchy, $v_k^{(b)}$, same drill as before: we ignore terms that do not involve $v_k^{(b)}$ for the particular node $b$ of interest in the objective (4). This time we collect terms involving the parent node of $b$, i.e. Pa$(b)$, and nodes $c$ whose parent is $b$, i.e. Pa$(c) = b$:

$$O = \alpha \left( \left\| v_k^{(b)} - v_k^{Pa(b)} \right\|_2^2 + \sum_{c \in \text{Child}(b)} \left\| v_k^{(c)} - v_k^{(b)} \right\|_2^2 \right) + C \tag{25}$$

$$= \alpha \left( v_k^{(b)} - v_k^{Pa(b)} \right)^\top \left( v_k^{(B)} - v_k^{Pa(b)} \right) + \alpha \sum_{c \in \text{Child}(b)} \left( v_k^{(c)} - v_k^{(b)} \right)^\top \left( v_k^{(c)} - v_k^{(b)} \right) + C \tag{26}$$

$$\begin{aligned} &= \alpha \left[ v_k^{(b)\top} v_k^{(b)} - 2 v_k^{(b)\top} v_k^{\text{Pa}(b)} + v_k^{\text{Pa}(b)\top} v_k^{\text{Pa}(b)} \right] \\ &\quad + \alpha \sum_{c \in \text{Child}(b)} \left[ v_k^{(c)\top} v_k^{(c)} - 2 v_k^{(c)\top} v_k^{(b)} + v_k^{(b)\top} v_k^{(b)} \right] + C \end{aligned} \tag{27}$$

$$= \alpha v_k^{(b)\top} v_k^{(b)} - 2\alpha v_k^{(b)\top} v_k^{\text{Pa}(b)} - \sum_{c \in \text{Child}(b)} 2\alpha v_k^{(c)\top} v_k^{(b)} + \sum_{c \in \text{Child}(b)} \alpha v_k^{(b)\top} v_k^{(b)} + C \tag{28}$$

Now we take the derivative, set to 0, and solve:

$$\frac{\partial O}{\partial v_k^{(b)}} = 2\alpha v_k^{(b)} - 2\alpha v_k^{\text{Pa}(b)} - \sum_{c \in \text{Child}(b)} 2\alpha v_k^{(c)} + \sum_{c \in \text{Child}(b)} 2\alpha v_k^{(b)} \tag{29}$$

$$0 = v_k^{(b)} - v_k^{\text{Pa}(b)} - \sum_{c \in \text{Child}(b)} v_k^{(c)} - |\text{Child}(b)| \cdot v_k^{(b)} \tag{30}$$

$$= (1 + |\text{Child}(b)|) v_k^{(b)} - v_k^{\text{Pa}(b)} - \sum_{c \in \text{Child}(b)} v_k^{(c)} \tag{31}$$

$$v_k^{(b)} = \frac{v_k^{\text{Pa}(b)} + \sum_{c \in \text{Child}(b)} v_k^{(c)}}{1 + |\text{Child}(b)|} \tag{32}$$

where $|\text{Child}(b)|$ is the number of direct child nodes of $b$.
